# Genetic and neural bases of the neuroticism general factor

**DOI:** 10.1101/2023.03.08.531776

**Authors:** Yuri Kim, Gretchen R. B. Saunders, Alexandros Giannelis, Emily A. Willoughby, Colin G. DeYoung, James J. Lee

## Abstract

We applied structural equation modeling to conduct a genome-wide association study (GWAS) of the general factor measured by a neuroticism questionnaire administered to *∼*380,000 participants in the UK Biobank. We categorized significant genetic variants as acting either through the neuroticism general factor, through other factors measured by the questionnaire, or through paths independent of any factor. Regardless of this categorization, however, significant variants tended to show concordant associations with all items. Bioinformatic analysis showed that the variants associated with the neuroticism general factor disproportionately lie near or within genes expressed in the brain. Enriched gene sets pointed to an underlying biological basis associated with brain development, synaptic function, and behaviors in mice indicative of fear and anxiety. Psychologists have long asked whether psychometric common factors are merely a convenient summary of correlated variables or reflect coherent causal entities with a partial biological basis, and our results provide some support for the latter interpretation. Further research is needed to determine the extent to which causes resembling common factors operate alongside other mechanisms to generate the correlational structure of personality.

## 1 Introduction

The biological underpinnings of personality are far from being understood. Genome-wide association studies (GWAS) can provide insight into personality’s biological etiology by indicating which genomic polymorphisms are significantly associated with a trait of interest. Most GWAS focus on single-nucleotide polymorphisms (SNPs), the most common type of genetic variation. SNPs reaching statistical significance in GWAS often lie near protein-coding genes and non-coding functional regions. As many functions of genes and their tissue-specific patterns of expression have been experimentally elucidated or computationally predicted, researchers can then infer the biological processes that are likely to be responsible for variation in the trait. Unfortunately, GWAS of personality traits often lack sample sizes large enough to detect many significant loci (e.g., Lo et al., 2017).

Studies focusing on neuroticism typically have been more successful (de Moor et al., 2015; Luciano et al., 2018; Nagel et al., 2018a; Okbay et al., 2016a; Smith et al., 2016). Neuroticism is one of the factors in the Big Five model of personality. Individuals who score highly in neuroticism tend to experience diverse and relatively more intense negative emotions. The largest GWAS meta-analysis of neuroticism to date found 136 significant independent loci (Nagel et al., 2018a). Neuroticism was measured using the Eysenck Personality Questionnaire–Revised Short Form (EPQ) (Eysenck et al., 1985). In the present study, we further investigated the genetics and biology of neuroticism using the summary statistics of a companion study analyzing the individual items in the questionnaire (Nagel et al., 2018b).

We also examined whether the significant SNPs act in accordance with the common-factor model, which is an important tool in the psychology of individual differences. McDonald (2003) suggested that a common factor might be regarded as a mental property with a non-physicalist interpretation, which nevertheless can be acted upon by physical causes: “the external variable causes the common factor of the dependent variables, that is, acts to change the level of the psychological attribute common to them” (p. 221). Others have proposed that a common-factor model is merely a convenient summary of otherwise formidably high-dimensional data rather than a representation or approximation of a causal model (Cramer et al., 2012). Genetics now provides us with an unprecedented opportunity to test these ideas. If we could find candidate causal variables, such as SNPs in the human genome, that exert effects on the questionnaire items proportional to their factor loadings, then we would have powerful evidence that the common factor does indeed mediate biological causes and therefore cannot be dismissed as an artifact. That is, if the loadings of certain dependent variables on their common factor were *λ*_1_, *λ*_2_, and so forth, then a SNP with effects on those variables of *βλ*_1_, *βλ*_2_, and so forth would strongly suggest that the SNP has on effect of *β* on *something* very much like the common factor.

Conversely, if the effects of the SNPs failed to accord with the factor loadings, this would suggest looking toward proposals such as “bonds” (Thomson, 1951) or network models (Cramer et al., 2012) for a superior causal model explaining the item covariation. Either way, identification of the biological mechanisms mediating the effects of the SNPs can provide insight into the nature of the higher-level objects in the hierarchy of explanation—whether those objects are common factors, “bonds,” networks, or something else entirely. A number of authors have previously tested a similar idea with general intelligence (*g*) (Cox et al., 2019; Kievit et al., 2012; Lee et al., 2019). Their results were consistent with brain size being one of multiple factors that affect a unitary *g*.

In this work we do not claim to resolve this issue conclusively. We claim merely that if we do find SNPs associated with all indicators to a degree corresponding roughly with their factor loadings, then we have evidence that common biological causes are one kind of mechanism contributing to the covariation “accounted for” by the common-factor model.

To conduct this analysis of the common factor neuroticism, we turned to Genomic SEM, a software tool for applying factor and path models to genetic data (Grotzinger et al., 2019). We classified the GWAS-identified SNPs as working either through the general factor, the group factors that happen to be present in this questionnaire, or none of the above (i.e., through “independent pathways”). It is the SNPs in the latter category that might call into question the appropriateness of the common-factor model at a deeper biological level. We then used the bioinformatic software tool DEPICT (Pers et al., 2015) in an attempt to identify the tissues and biological mechanisms mediating the effects of the SNPs in these categories. In this way we not only tested the verisimilitude of the common-factor model at the genetic level, but also obtained mechanistic insight into the nature of the neuroticism factor. Eysenck (1992) in particular stressed the importance of grounding the constructs of personality models genetically and biologically in order to further their validity.

## 2 Methods

### 2.1 Confirmatory factor analysis

We used the software tool Genomic SEM (Grotzinger et al., 2019) to calculate the genetic covariance matrix of the neuroticism items in the Eysenck Personality Questionnaire–Revised Short Form, as administered to about 380,000 UK Biobank participants (Nagel et al., 2018b). The “genetic correlation” between two traits is the correlation between their heritable components. That is, if each trait is the sum of a genetic and environmental term, then the genetic correlation is the correlation between just the genetic terms. Genetic correlations tend to be close to their corresponding phenotypic correlations (Sodini et al., 2018), being slightly larger on average, and so should yield a similar factor-analytic solution (e.g., de la Fuente et al., 2021). To calculate the genetic correlation between two binary traits, estimates of the population prevalences (pass rates) are required. We used the estimates previously published (Nagel et al., 2018b). Note that the genetic correlations are calculated over essentially all “common SNPs”—polymorphic sites where both alleles exceed a threshold frequency—regardless of statistical significance.

We adopted the three-factor model of the neuroticism questionnaire used in the original Genomic SEM publication by Grotzinger et al. (2019). In this model the items *mood*, *misery*, *irritable*, *fed-up*, and *lonely* are indicators of a factor that we will call depressed affect, after the largely similar group of items identified by hierarchical cluster analysis (Nagel et al., 2018b). The items *nervous*, *worry*, *tense*, and *nerves* are indicators of a factor that we will call worry, also after a similar cluster identified in the previous analysis. The group factors depressed affect and worry do not readily map onto aspects in the BFAS (DeYoung et al., 2007), but do arguably map onto the respective facets depression and anxiety in the NEO (Costa & McCrae, 1992). The items *guilt*, *hurt*, and *embarrass* are indicators of a third factor that we will call vulnerability, after the largely similar group of items identified by exploratory factor analysis (Hill et al., 2020). We introduced a neuroticism general factor into this model by treating the three group factors as indicators of a hierarchical second-order factor. Unit-variance identification was employed.

There is some evidence that participants in the UK Biobank tend to be slightly less neurotic than the rest of the population (Tyrrell et al., 2021; Young et al., 2022). Such selection bias can distort the factor structure of the measurements (Lee, 2012; Meredith, 1993). Our conjecture is that psychological traits most affecting participation in research are those related to education and social class, and neuroticism does not seem strongly related to such status markers (Demange et al., 2021; Mammadov, 2022; Poropat, 2009; Zell & Lesick, 2022). When the association between personality and research participation has been directly studied, no significant correlations with neuroticism have been observed (Cheng et al., 2020; Marcus & Schütz, 2005). Therefore we expect any impact of selection bias on our results to be modest.

### 2.2 Path modeling of SNP effects

#### 2.2.1 GWAS of the neuroticism general factor

We performed a GWAS of the neuroticism general factor by specifying, in Genomic SEM, a path from the tested SNP to the second-order general factor (Fig. 1A). Any confounding with non-genetic variables is likely to be minimal because within-family GWAS of the neuroticism sum score have produced results very close to those of population GWAS (Howe et al., 2022; Young et al., 2022). We used the reference file supplied by Genomic SEM to retain only SNPs with a minor allele frequency (MAF) exceeding .005 in the 1000 Genomes European populations. This left more than 7 million SNPs in the GWAS. Additional methodological details of both the original item-level GWAS and our GWAS at the latent level with Genomic SEM are given in the Supplementary Material.

**Figure 1:**
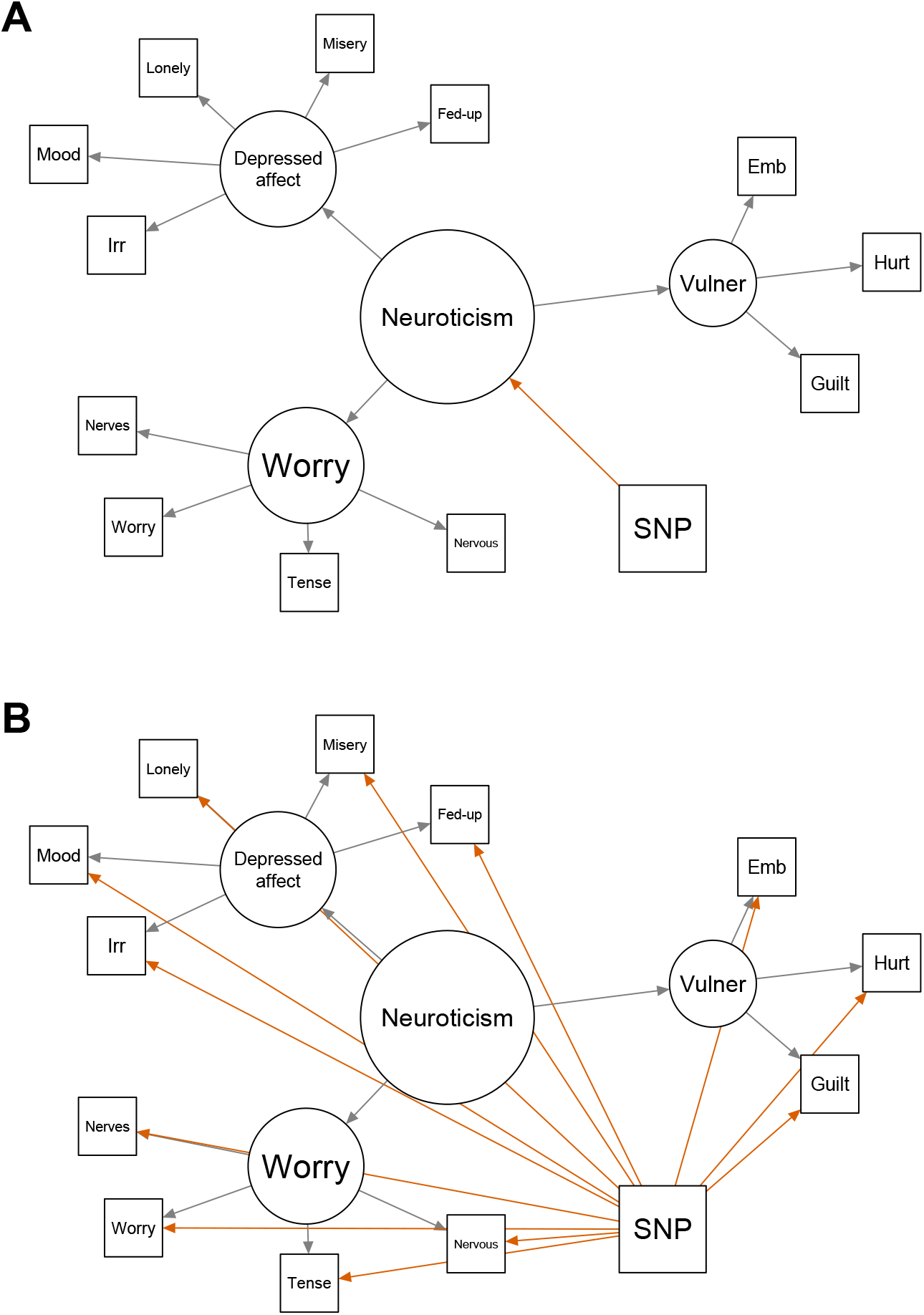
Path diagrams portraying how a single-nucleotide polymorphism (SNP) might be associated with the questionnaire items. A. The focal SNP (or a nearby highly correlated SNP) acts through the neuroticism general factor. B. The focal SNP (or a nearby highly correlated SNP) acts on the 12 items through “independent pathways.” Not shown is a model where the SNP’s associations are with one or more of the three group factors.

Because they are often highly correlated, nearby SNPs may not not represent independent association signals. We attempted to identify independently significant SNPs by using the “clump” function of the software tool PLINK (Chang et al., 2015; Purcell et al., 2007). In essence, clumping picks out local minima of the *p*-value sequence along the genome. We used the clump settings of the bioinformatics tool DEPICT (Pers et al., 2015), which calls PLINK to identify lead SNPs. The most important of these settings is the threshold *p <* 10*^−^*^5^ for the statistical significance of the association between SNP and trait. Although less stringent than the conventional GWAS significance threshold *p <* 5 *×* 10*^−^*^8^, this threshold is recommended by the DEPICT developers because the biological annotation provided by their tool (see below) is tolerant of false-positive SNPs.

Note that the conventional GWAS threshold aspires to prevent even a single false positive from appearing among the SNPs significantly associated with a single trait. Although there may be at least one false positive among the SNPs in the range 10*^−^*^5^ *> p ≥* 5 *×* 10*^−^*^8^, many of these SNPs will be true positives in a well-powered GWAS with many SNPs reaching *p <* 5 *×* 10*^−^*^8^.

We subjected the candidate lead SNPs from the GWAS of the neuroticism general factor to further tests. We ran a “group-factor” model in which the three first-order group factors were regressed on each of the candidate lead SNPs. This model thus requires three path coefficients in the place of the one required by the general-factor model. The general-factor model is nested within the group-factor model, the former being obtained from the latter by making the three SNP effects proportional to the loadings of the group factors on the general factor. We then ran an “independent-pathway” model regressing all 12 items on each candidate lead SNP (Fig. 1B). The independent-pathway model thus estimates 12 path coefficients in the place of the three required by the group-factor model; the latter is nested within the former.

The independent-pathway model is an operationalization of not only Thomson’s bonds model, but also the network model (Cramer et al., 2012); our Fig. 1 contrasting the common-factor and independent-pathway models is exactly parallel to Figure 7 of Cramer et al. (2012). These authors proposed that support for the independent-pathway model over the common-factor model would count as support for their network perspective. Taking the most significant SNPs in the GWAS of neuroticism sum scores published at that time, they carried out an analysis similar to ours and claimed to find some evidence for the SNPs acting on individual items rather than the general factor. The only SNP-item association of theirs that we could attempt to look up and replicate was the one between rs12509930 and *guilt*. In the UK Biobank sample of roughly 380,000 individuals, this association is not significant (*p* = .70). We should not be surprised by this replication failure, in light of the small sample sizes of the GWAS at that time, and the authors themselves avowed the tentative and exploratory nature of their analysis. The important point is that we can now carry out their proposal of pitting the common-factor and network models against each other to a much greater extent than was possible a decade ago.

To determine whether a candidate lead SNP identified in the GWAS of the neuroticism general factor is better regarded as acting through factors or independent pathways, one can test the significance of the difference in *χ*^2^ between more and less parsimonious models. The Genomic SEM developers call this difference *Q*_SNP_ (Genomic SEM tutorial, accessed October 2020). In one of their analyses, Grotzinger et al. (2019) used the threshold *p > .*005 for calling a *Q*_SNP_ value “low.” Following the suggestion of a reviewer, however, we carried out model selection using Akaike weights (Wagenmakers & Farrell, 2004). The sum of the weights equals one by construction, making them analogous to probabilities. The ratio of two weights can be interpreted as the relative likelihood of the model corresponding to the numerator (Royall, 1997) times a factor penalizing that model if it has more estimated parameters. Such a penalty may be desirable if a sufficient increase in sample size will lead to the rejection of any simple model regardless of its qualitatively excellent fit. We treated any model with an Akaike weight exceeding 2*/*3 as the “correct” model for a given SNP, as this means at least twice as much support as any alternative. It is possible for no model to obtain this large a weight, meaning that the SNP’s associations with the items are not clearly fit best by any of the candidate models.

Since calculating the model *χ*^2^ and AIC increased the computation time of a SNP association by roughly a factor of 10 in the version of Genomic SEM that we used (October 2020), we did not calculate these for all SNPs in the GWAS but rather only the lead SNPs, once for each of the three candidate models (general factor, group factor, independent pathway). Supplementary Fig. S1 provides an overview of our pipeline for the GWAS of the neuroticism general factor and subsequent classification of lead SNPs.

#### 2.2.2 GWAS of additional factors

We also conducted GWAS of each group factor with nontrivial variance attributable to sources other than the neuroticism general factor (i.e., depressed affect and worry). The first step of our procedure was to conduct a GWAS with Genomic SEM, specifying directed edges from the SNP to all three group factors. We then examined each factor’s association results satisfying *p <* 10*^−^*^5^. Of the lead SNPs identified by the clumping procedure, we discarded any already assigned to either the general-factor or independent-pathway model in the GWAS of the neuroticism general factor (Supplementary Fig. S1). Since we were particularly interested in SNPs associated solely with the focal group factor, we tested each remaining lead SNP for association with that factor while setting to zero the coefficients of its paths to the other two factors. We also ran the independent-pathway model for each of these lead SNPs (Fig. 1B). As before, we used an Akaike weight exceeding 2*/*3 as the criterion for assigning a lead SNP to one of three competing models (all group factors, one group factor, independent pathways). Supplementary Fig. S2 provides an overview of our pipeline for the GWAS of the group factors and subsequent classification of lead SNPs.

To convey the difference between this GWAS and the one outlined in Supplementary Fig. S1, we will give an example of a SNP that would be ascertained as significant in the former but not in the latter. Suppose that a SNP acts solely through the residual of a group factor. This SNP might be ascertained in the GWAS of the group factors, through a combination of a relatively large effect size and favorable sampling variation. It might not be ascertained in the GWAS of the general factor, despite this GWAS containing a follow-up step checking for association with the group factors, because it is less likely to become a lead SNP in the first step. This difference in the ascertainment scheme can be important for certain inferences, a matter to which we return in the Discussion.

It is worthwhile to consider whether independent-pathway SNPs enrich any tissues or biological pathways (see below), despite not acting through any common factors. To identify such SNPs, Grotzinger et al. (2019) conducted two GWAS, one of neuroticism in their single-factor model and the other of independent pathways, and calculated a form of the *Q*_SNP_ statistic for each SNP in the GWAS. At the time of our own analysis, this procedure was beyond the computational resources available to us. As a compromise, we took forward to DEPICT the union of the lead SNPs from the GWAS of the common factors that qualified by virtue of their Akaike weights for the independent-pathway model.

### 2.3 Genetic correlations

Genomic SEM calls LD Score regression (LDSC) to calculate genetic correlations, and this method is known to be unbiased under fairly general conditions (Bulik-Sullivan et al., 2015; Lee et al., 2018a).

A finding of genetic correlations similar to those calculated in previous studies of neuroticism observed scores would provide an affirmative quality-control check of our approach based on structural equation modeling. It would also support the validity of the common assumption that a correlation with an observed sum score primarily reflects a correlation with the scale’s general factor. The Supplementary Material lists the traits used in this analysis and accompanying references.

We also calculated genetic correlations with the residuals of the group factors depressed affect and worry. Procedurally we used Genomic SEM to specify the bifactor model generalizing the hierarchical model displayed in Fig. 1 and then performed a GWAS of the group factors within the bifactor model. Supplementary Fig. S3 displays the factor and path model that we employed for this purpose. We used the resulting GWAS summary statistics to calculate the genetic correlations with depressed affect and worry.

Supplementary Fig. S4 and Supplementary Table S1 present the results.

### 2.4 Polygenic prediction

At the request of a reviewer, we used the summary statistics from our GWAS of the common factors to calculate polygenic scores (PGS) and validate them in a new sample. Methodological details are given in the Supplementary Material, and Supplementary Table S2 presents the results.

### 2.5 Biological annotation

#### 2.5.1 DEPICT

DEPICT (Data-driven Expression Prioritized Integration for Complex Traits) is a software tool that prioritizes likely causal genes affecting the trait, identifies tissues/cell types where the causal genes are highly expressed, and detects enrichment of gene sets. A “gene set” is a group of genes designated by database curators as sharing some common property, such as encoding proteins that participate in the same biological function. A gene set shows “enrichment” if SNPs significantly associated with the trait fall in or near the set’s member genes more often than expected by chance. More complete descriptions of DEPICT can be found in previous publications (Okbay et al., 2016b; Pers et al., 2015).

Our path modeling with Genomic SEM placed each lead SNP into a collection (e.g., SNPs associated with the neuroticism general factor). Each such collection of SNPs was supplied as input to DEPICT (https://github.com/perslab/DEPICT, release 194). DEPICT takes lead SNPs and merges them into loci potentially encompassing more than one lead SNP according to certain criteria (Pers et al., 2015). The genes overlapping these loci are the basis of the DEPICT analysis. The limitation of the DEPICT input to a subset of SNPs is an important strength in our application. A method that relies on genome-wide summary statistics is not straightforward to adapt if some SNPs in a GWAS of a common factor must be dropped for better fitting a more complex model (Fig. 1).

To run DEPICT, we edited and then executed the template configuration file. We left in place all default parameter values except those affecting how the results are printed in the output files. We also used a collections file of the genes overlapping the locus around a given SNP based on 1000 Genomes phase 3 rather than 1000 Genomes pilot data; this file was given to us by the DEPICT developers and is available along with the GWAS summary statistics generated for this study. Many tissues/cell types and gene sets in the DEPICT inventory are in fact duplicates despite having different identifiers; we adopted the pruned list of tissues/cell types used by Finucane et al. (2018) and excluded duplicated gene sets using the criteria set out by Lee et al. (2018b). Except where noted, we adopted the developer-recommended definition of statistical significance at the level of genes, tissues/cell types, and gene sets as a false discovery rate (FDR) below .05.

The reconstitution of the gene sets was motivated by a desire to compensate for the limitations of existing bioinformatic databases, which suffer from both false positives and false negatives. The reader can consult Supplementary Table 28 of Lee et al. (2018b) for a demonstration of the reconstitution procedure’s success in empowering detection of enrichment only in sets appropriate to the studied trait. The reconstitution procedure has also proven fruitful in other applications (Cvejic et al., 2013; Fehrmann et al., 2015).

#### 2.5.2 Stratified LD Score regression and PANTHER overrepresentation test

At the request of a reviewer, we have calculated effect sizes in terms of fold enrichment to accompany the displays of statistically significant results in Figure 2 and Table 2. We used two different tools for this purpose. The first was stratified LD Score regression (S-LDSC), a standard method for testing enrichment of discrete gene sets (Finucane et al., 2015). The enrichment statistic calculated by S-LDSC is

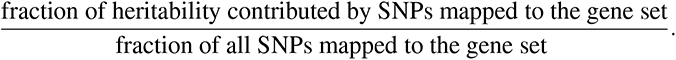

**Figure 2:**
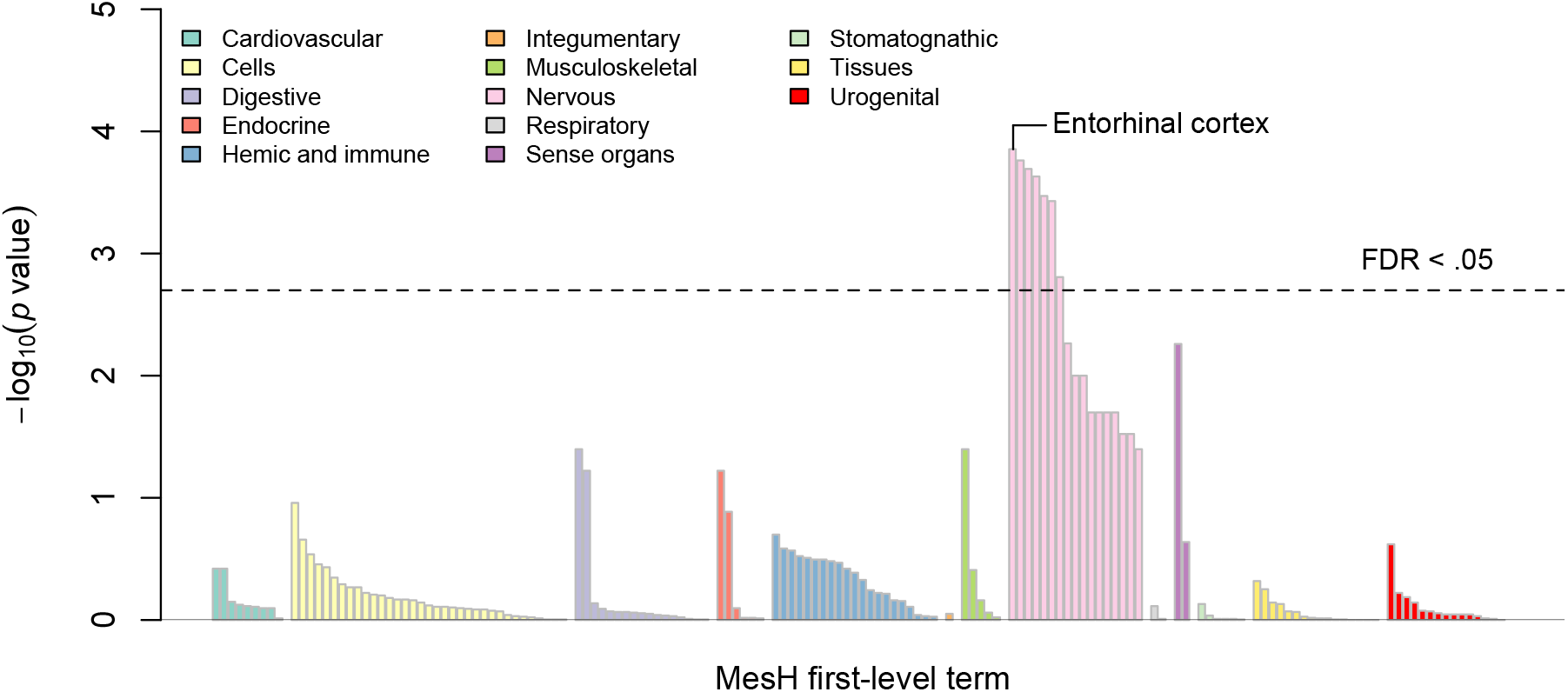
Tissues or cell types with significant expression of genes near SNPs associated with the neuroticism general factor (relative to genes in random sets of loci). The tissues are arranged along the *x*-axis by Medical Subject Heading (MeSH) first-level term. The *y*-axis represents statistical significance on a *−* log_10_ scale. The height of the dashed horizontal line corresponds to the *p* value yielding FDR *< .*05. See Supplementary Table S6 for complete results.

“Gene set” here can equally well mean a group of genes that are highly expressed in a given tissue/cell type. We employed the Finucane et al. (2018) procedure of taking the top 10 percent of genes in the DEPICT inventory belonging to a given gene set, mapping all SNPs lying within 100 kb of a member gene to that set, and using the so-called baseline annotations and an any-gene indicator as control variables. We used the 97 baseline annotations currently recommended by the developers (downloaded August 2023 from https://storage.googleapis.com/broad-alkesgroup-public-requester-pays/LDSCORE). We also used the precomputed stratified LD Scores for the DEPICT tissues/cell types supplied by the developers (“Franke dataset”). The developers state that they provide a gene-coordinate file so that users can calculate their own stratified LD Scores for novel gene sets (LD Score estimation tutorial, accessed August 2023), but this file seems not to have been transferred to their Google Cloud depository. To calculate stratified LD Scores for the reconstituted gene sets found to be significantly enriched in the standard DEPICT analysis, we used instead the latest version of the GENCODE coordinate file (downloaded August 2023), taking the row in this file assuming the value of *gene* in the *feature* column as providing the canonical start and stop coordinates of a given Ensembl identifier. The standard 1-centimorgan radius was used to calculate the stratified LD Scores.

We tested the null hypothesis that the enrichment is equal to one. Previous experience with this method suggests that a 1.3-fold enrichment of a gene set should be regarded as a large effect size (Finucane et al., 2018; Kim et al., 2019; Lee et al., 2018b), although smaller non-null sets and sets specifically constructed to contain genes under strong purifying selection may yield higher values.

Our second method for calculating fold enrichments was the PANTHER overrepresentation test, which has been implemented as a web-based tool (http://www.geneontology.org). The input to this method is a discrete list of genes supplied by the user. To increase statistical power, we used the Ensembl identifiers of all DEPICT-prioritized genes satisfying FDR *< .*20 as input. Standard FDR calculations assume that the alternative hypothesis is true in only a small proportion of cases, and a violation of this assumption leads to the FDR being conservative (Efron, 2010). As there is almost certainly a causal gene near most lead SNPs, many genes falling in the interval .05 *≤* FDR *< .*20 are likely to be true positives. We used all default settings for analyses launched from the front page of the Gene Ontology website.

The null hypothesis in the PANTHER overrepresentation test is that the input gene list is a random sample of all genes in the reference gene list. The enrichment statistic is thus

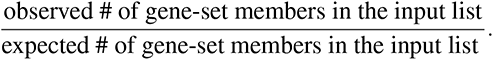

The PANTHER overrepresentation test has properties that complement those of S-LDSC. It is based on the discrete version of the gene set rather than the reconstituted version and thus provides a way to check the robustness of the latter. (The PANTHER database does not include the Mammalian Phenotype gene sets from the Mouse Genomics Institute.) Furthermore, it is arguably testing a hypothesis that is closer to the one being tested by the standard DEPICT analysis. In the latter approach, we are asking whether the lead SNPs at the current stage of a GWAS fall disproportionately within or near high-ranking members of a given gene set. The answer to this question may change as the GWAS increases in sample size and begins to add different types of SNPs. In contrast, S-LDSC is calculating a measure of enrichment that applies to the whole genome rather than a subset of SNPs. In theory, the S-LDSC enrichment statistic does not change as the GWAS progresses, although the standard error of its estimate hopefully grows smaller. The PANTHER overrepresentation test is closer in spirit to the standard DEPICT approach in that it focuses on genes that happen to encompass or lie near the current lead SNPs.

## 3 Results

### 3.1 Factor analysis of the neuroticism questionnaire

We replicated the indices reported by Grotzinger et al. (2019) indicating a good fit of a model with three group factors (CFI = .969, SRMR = .054). We therefore regarded the three-factor model as satisfactory for purposes of SNP-level path modeling. The loading of the vulnerability group factor defined by *guilt*, *hurt*, and *embarrass* on the neuroticism general factor was estimated to be nearly one (.97) (Supplementary Table S3). These items seem to have very little genetic variance shared in common other than what is attributable to neuroticism. For this reason we did not conduct a GWAS of this factor when trying to identify SNPs associated with group factors. Although our result here may seem to diverge from that of Hill et al. (2020), their bifactor model allowed correlations between group factors and thus qualitatively differed from our hierarchical model. As discussed in the Supplementary Material, we did by and large replicate the Hill et al. (2020) finding of markedly different genetic correlations of the neuroticism general factor and the residual worry factor with certain traits (Supplementary Fig. S4).

### 3.2 GWAS of the neuroticism general factor

Before examining the main results and downstream analyses of a GWAS, it is reasonable to assess the overall amount of signal present in its summary statistics. The product of the sample size and the heritability (e.g., as estimated by LD Score regression) is normally a good metric for this purpose, but it is inapplicable to a GWAS of a latent trait conducted with Genomic SEM because neither factor in this product is well defined (Mallard et al., 2022). We followed the recommendation of the Genomic SEM developers to use the mean *χ*^2^ statistic instead (Supplementary Table S4). The mean *χ*^2^ of our neuroticism GWAS was 1.63—very close to those of past groundbreaking GWAS of behavioral traits (Okbay et al., 2016b; Pers et al., 2016; Schizophrenia Working Group of the Psychiatric Genomics Consortium, 2014). Our GWAS summary statistics seem to contain sufficient signal for meaningful downstream analyses. Note that an undefined heritability is not a problem in the use of LDSC to obtain genetic correlations and functional enrichments because of cancellations from numerator and denominator in the calculations of those quantities.

Our GWAS of the neuroticism general factor identified 394 lead SNPs satisfying *p <* 10*^−^*^5^, in 296 distinct DEPICT-defined loci. We examined these SNPs for an improvement in model fit upon increasing the number of paths. Thirty-five of the 394 SNPs were characterized by small negative values of the *Q*_SNP_ statistic when comparing the fit of the model where the SNP acts on the general factor (Fig. 1A) to that of the model where the SNP acts on the three group factors. Such negative values can arise when the two models under comparison are distinguished by few degrees of freedom, and they indicate an extremely good fit of the data to the more restrictive model (A. Grotzinger, personal communication). Of the 394 lead SNPs, 139 qualified by virtue of their Akaike weights for the general-factor model, 81 for the group-factor model, and 63 for the independent-pathway model. One hundred eleven SNPs had no Akaike weight greater than 2*/*3, precluding for now their assignment to any model. Of these 111 indeterminate SNPs, a plurality of 54 attained their largest Akaike weight in the general-factor model.

Supplementary Table S5. lists the 139 general-factor lead SNPs. Nineteen of these SNPs attained the strict genome-wide significance level *p <* 5 *×* 10*^−^*^8^ (Table 1). Of these 19 SNPs, 17 reached strict genome-wide significance in the largest GWAS to date of an observed neuroticism score (Nagel et al., 2018a). Information about all significant SNPs regardless of classification can be found in the Supplementary Data.

**Table 1:**
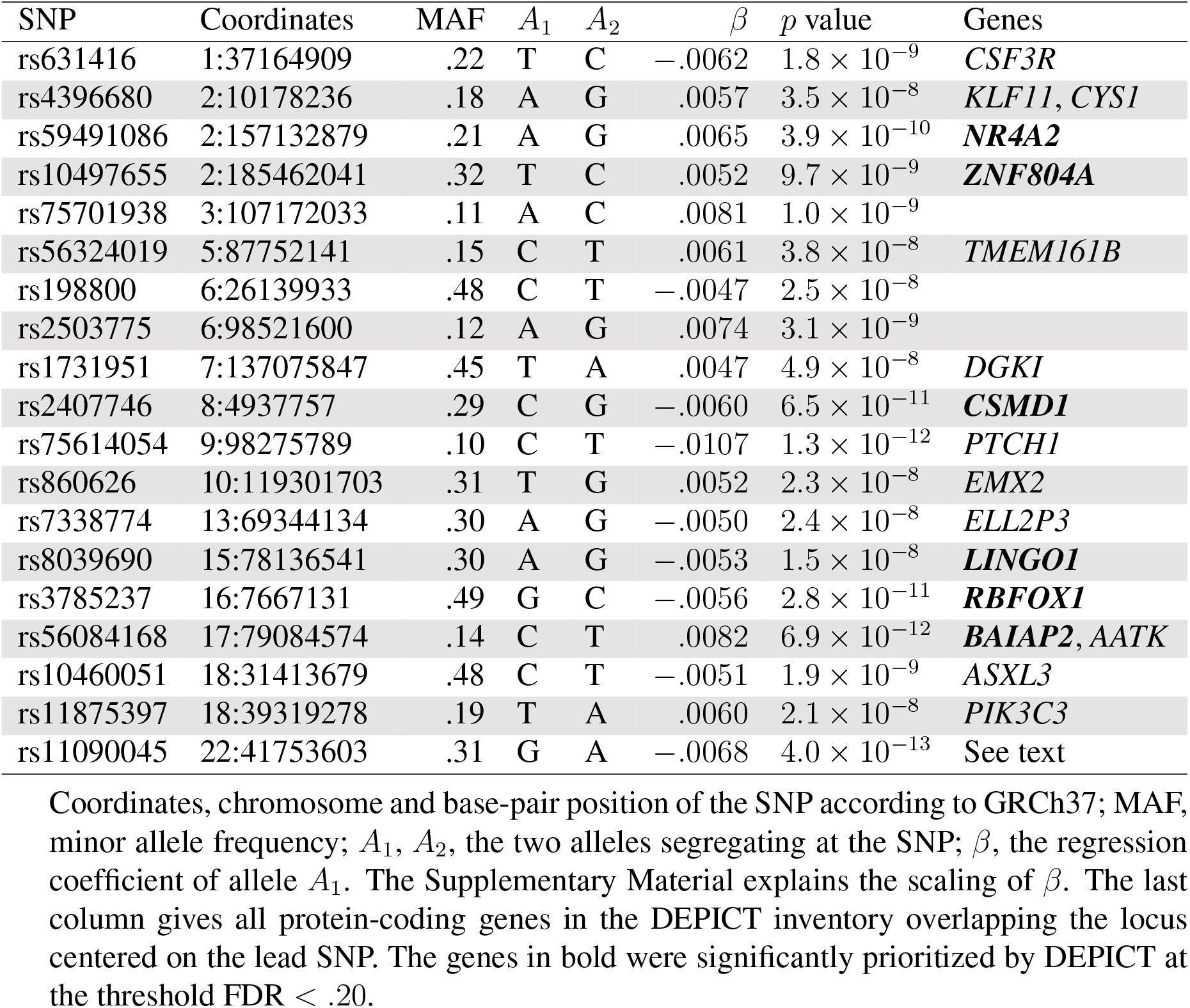
Strictly genome-wide significant SNPs in the GWAS of the neuroticism general factor with Akaike weight *>* 2*/*3 for the model in Fig. 1A.

| SNP | Coordinates | MAF | $A_1$ | $A_2$ | $\beta$ | $p$ value | Genes |
| --- | --- | --- | --- | --- | --- | --- | --- |
| rs631416 | 1:37164909 | .22 | T | C | -.0062 | $1.8 \times 10^{-9}$ | *CSF3R* |
| rs4396680 | 2:10178236 | .18 | A | G | .0057 | $3.5 \times 10^{-8}$ | *KLF11, CYS1* |
| rs59491086 | 2:157132879 | .21 | A | G | .0065 | $3.9 \times 10^{-10}$ | ***NR4A2*** |
| rs10497655 | 2:185462041 | .32 | T | C | .0052 | $9.7 \times 10^{-9}$ | ***ZNF804A*** |
| rs75701938 | 3:107172033 | .11 | A | C | .0081 | $1.0 \times 10^{-9}$ | |
| rs56324019 | 5:87752141 | .15 | C | T | .0061 | $3.8 \times 10^{-8}$ | *TMEM161B* |
| rs198800 | 6:26139933 | .48 | C | T | -.0047 | $2.5 \times 10^{-8}$ | |
| rs2503775 | 6:98521600 | .12 | A | G | .0074 | $3.1 \times 10^{-9}$ | |
| rs1731951 | 7:137075847 | .45 | T | A | .0047 | $4.9 \times 10^{-8}$ | *DGKI* |
| rs2407746 | 8:4937757 | .29 | C | G | -.0060 | $6.5 \times 10^{-11}$ | ***CSMD1*** |
| rs75614054 | 9:98275789 | .10 | C | T | -.0107 | $1.3 \times 10^{-12}$ | *PTCH1* |
| rs860626 | 10:119301703 | .31 | T | G | .0052 | $2.3 \times 10^{-8}$ | *EMX2* |
| rs7338774 | 13:69344134 | .30 | A | G | -.0050 | $2.4 \times 10^{-8}$ | *ELL2P3* |
| rs8039690 | 15:78136541 | .30 | A | G | -.0053 | $1.5 \times 10^{-8}$ | ***LINGO1*** |
| rs3785237 | 16:7667131 | .49 | G | C | -.0056 | $2.8 \times 10^{-11}$ | ***RBFOX1*** |
| rs56084168 | 17:79084574 | .14 | C | T | .0082 | $6.9 \times 10^{-12}$ | ***BAIAP2, AATK*** |
| rs10460051 | 18:31413679 | .48 | C | T | -.0051 | $1.9 \times 10^{-9}$ | *ASXL3* |
| rs11875397 | 18:39319278 | .19 | T | A | .0060 | $2.1 \times 10^{-8}$ | *PIK3C3* |
| rs11090045 | 22:41753603 | .31 | G | A | -.0068 | $4.0 \times 10^{-13}$ | See text |

Coordinates, chromosome and base-pair position of the SNP according to GRCh37; MAF, minor allele frequency; $A_1$ , $A_2$ , the two alleles segregating at the SNP; $\beta$ , the regression coefficient of allele $A_1$ . The Supplementary Material explains the scaling of $\beta$ . The last column gives all protein-coding genes in the DEPICT inventory overlapping the locus centered on the lead SNP. The genes in bold were significantly prioritized by DEPICT at the threshold $\text{FDR} < .20$ .

The most significant general-factor SNP was rs11090045 (*p* = 4.0 *×* 10*^−^*^13^). Its locus on chromosome 22 is a very gene-dense region, overlapping *ZC3H7B* (FDR *< .*05), *TEF* (FDR *< .*20), *TOB2* (FDR *< .*20), *CSDC2* (FDR *< .*20), *EP300* (FDR *< .*20), *PMM1*, *RANGAP1*, *XRCC6*, *CHADL*, *ACO2*, *L3MBTL2*, *PPDE2*, *PHF5A*, and *POLR3H*. Although rs11090045 itself is located in the 3*^′^* untranslated region of *ZC3H7B*, the unusual number of candidates for causal genes in this locus may possibly be explained by the hypothesis of rs11090045 being a correlated proxy for multiple causal SNPs collectively acting through more than one gene.

It is of interest to examine how the cutoffs defined by Akaike weights correspond to *Q*_SNP_ statistics. Upon treating any SNP with a negative *Q*_SNP_ statistic as having a *p* value of one, we found that the 139 SNPs assigned by their Akaike weights to the general-factor model were all characterized by *p > .*28 (median *p* = .68) with respect to the null hypothesis of the general-factor model fitting better than the group-factor model. If we take the *p < .*05 criterion as standard, then our use of Akaike weights to define general-factor SNPs seems conservative. In contrast, for the 63 SNPs qualifying for the independent-pathway model, the *Q*_SNP_ *p* values with respect to the null hypothesis of the group-factor model fitting better than the independent-pathway model all met *p < .*02 (median *p* = .001).

### 3.3 Significant tissues/cell types and gene sets

The output of DEPICT provides insight into the biology associated with the SNPs appearing to act through the neuroticism general factor. Fig. 2 shows that there were 7 statistically significant tissues/cell types. All of these without exception bore the MeSH second-level term *central nervous system*. The most significant result was *entorhinal cortex* (*p* = 1.4 *×* 10*^−^*^4^). The entorhinal cortex is a way station connecting the neocortex, the hippocampus, and the amygdala, passing along signals critical for memory formation, navigation, and the perception of time (Maass et al., 2015; Tsao et al., 2018). The second most significant result was *limbic system* (*p* = 1.7 *×* 10*^−^*^4^), which refers to a collection of structures immediately below the medial temporal lobe that includes the entorhinal cortex and hippocampus. Overall, the neuroticism general factor showed the clear signature of a behavioral trait mediated by the brain.

More revealing than these tissue-level results were the significantly enriched gene sets. There were 21 such sets, and Table 2 shows the 6 of these that are not protein-protein interaction (PPI) subnetworks. *Abnormal cued conditioning behavior* (*p* = 6 *×* 10*^−^*^6^), *increased anxiety-related response* (*p* = 8.9 *×* 10*^−^*^5^), and *decreased exploration in new environment* (*p* = 9.1 *×* 10*^−^*^5^) are all taken from the Mouse Genome Informatics database and defined by fearful and anxious behavior when their member genes are perturbed in mice.

**Table 2:**
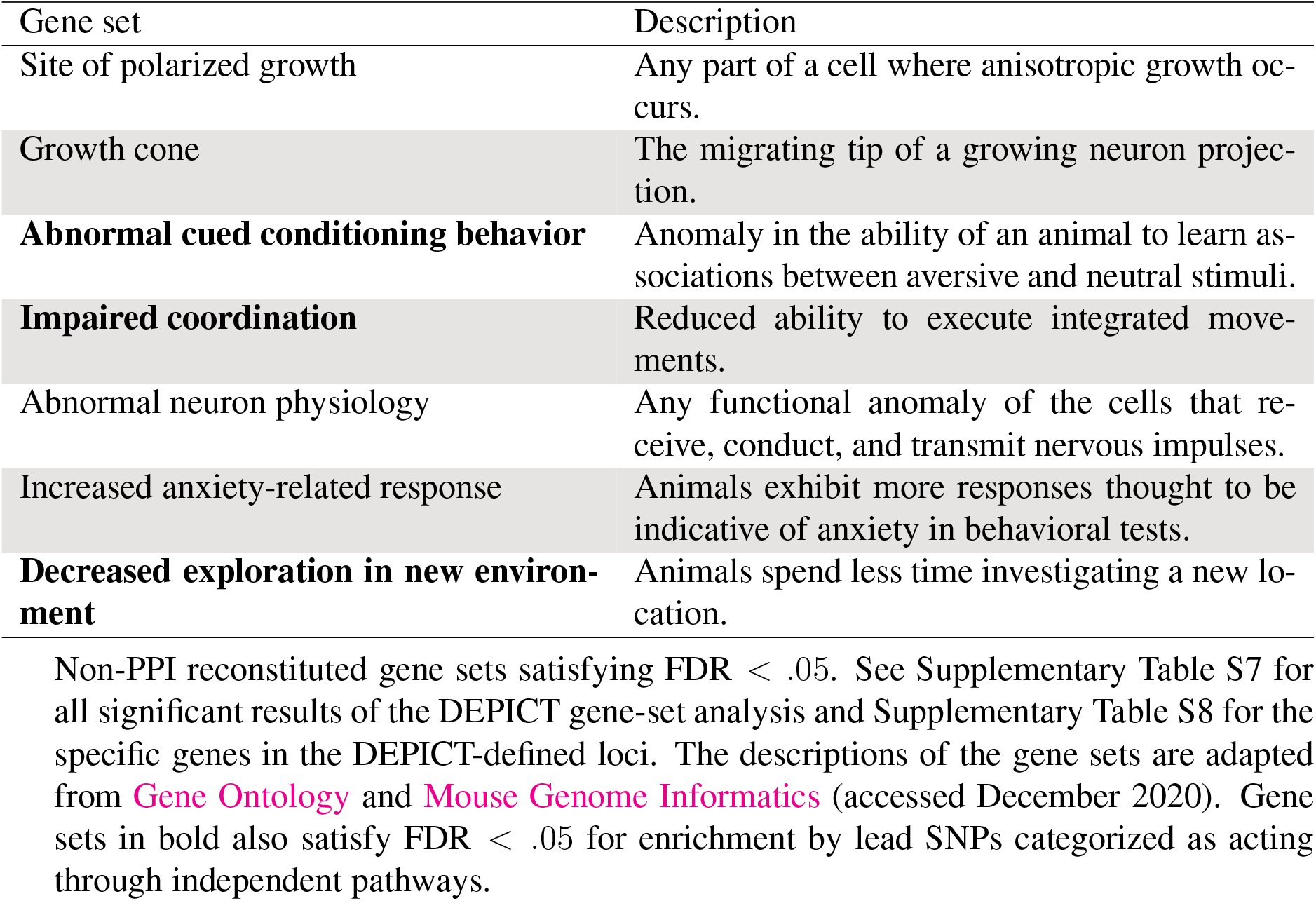
Reconstituted gene sets significantly enriched by lead SNPs for the neuroticism general factor.

| Gene set | Description |
| --- | --- |
| Site of polarized growth | Any part of a cell where anisotropic growth occurs. |
| Growth cone | The migrating tip of a growing neuron projection. |
| **Abnormal cued conditioning behavior** | Anomaly in the ability of an animal to learn associations between aversive and neutral stimuli. |
| **Impaired coordination** | Reduced ability to execute integrated movements. |
| Abnormal neuron physiology | Any functional anomaly of the cells that receive, conduct, and transmit nervous impulses. |
| Increased anxiety-related response | Animals exhibit more responses thought to be indicative of anxiety in behavioral tests. |
| **Decreased exploration in new environment** | Animals spend less time investigating a new location. |

### 3.4 GWAS of the group factors

We now report our attempts to find SNPs associated with the group factor depressed affect. Recall that we conducted a GWAS with Genomic SEM, based on a model sending directed edges from the SNP to all three group factors. After discarding SNPs identified as general-factor or independent-pathway SNPs in previous analyses, we ended up with 317 lead SNPs. Of these 317, 53 reached the strict genome-wide significance threshold *p <* 5 *×* 10*^−^*^8^. Interestingly, only 7 of the 317 lead SNPs were selected by the criterion of an Akaike weight greater than 2*/*3 as having no associations with the other two group factors, and none of these 7 reached the stringent genome-wide significance threshold *p <* 5 *×* 10*^−^*^8^. In contrast, 184 SNPs qualified by virtue of their Akaike weights for the group-factor model (nonzero effects on all three factors), 64 for the independent-pathway model, and 62 for none of the above.

The 184 SNPs qualifying for the group-factor model showed highly concordant effects on the three factors. In other words, despite being deemed a poor fit to the general-factor model, a SNP’s association with one factor was highly predictive of its associations with the two others. The sign concordance between SNP effects on depressed affect and worry was 100 percent. Each sign concordance between a major group factor and the vulnerability factor (with little non-neuroticism genetic variance) was 183/184.

After running the analogous procedure, we identified 286 lead SNPs associated with worry. Of these 286, 14 reached *p <* 5 *×* 10*^−^*^8^. Only 4 of the 286 lead SNPs were associated solely with the residual group factor of worry, none of which attained *p <* 5 *×* 10*^−^*^8^. Of the remaining SNPs, 184 qualified by virtue of their Akaike weights for the group-factor model, 54 for the independent-pathway model, and 43 for none of the above. The sign concordances were again either 100 percent or short of perfect by one SNP.

Supplementary Table S9. lists the 11 total SNPs associated with the residual group factors. Such a small number of lead SNPs, particularly when few reach strict genome-wide significance, leads to low statistical power with DEPICT (Turley et al., 2018). Therefore we did not conduct biological annotation of these 11 SNPs.

The Supplementary Data contain information about all of the SNPs used in these analyses.

### 3.5 Independent-pathway SNPs

Our analyses of the common factors assigned a total of 181 lead SNPs to the independent-pathway model (Supplementary Table S10), and we proceeded to annotate these. The significantly enriched tissues/cell types were, as expected, those of the nervous system, including *limbic system* (*p* = 4.7 *×* 10*^−^*^4^) and *entorhinal cortex* (*p* = 5.5 *×* 10*^−^*^4^) (Supplementary Table S11).

There were 27 significantly enriched gene sets (Supplementary Table S12). As indicated in Table 2, many were shared with the neuroticism general factor (*abnormal cued conditioning behavior*, *impaired coordination*, *decreased exploration in new environment*). One of the independent-pathway gene sets, *abnormal contextual conditioning behavior*, is also defined by the learning of fear and caution. The Mouse Genome Informatics database describes the relevant phenotype as an “anomaly in the ability of an animal to learn and remember an association between an aversive experience . . . and the neutral, unchanging environment” (accessed March 2023).

The other significant results pointed to the early development of the brain (e.g., *central nervous system neuron axonogenesis*) and synaptic activity in the behaving organism (e.g., *glutamatergic synaptic transmission*).

The SNPs were grouped into 112 loci that in turn overlapped 324 genes (Supplementary Table S13). Thirty of these 324 genes were also among the 228 genes overlapping the loci encompassing the lead SNPs for the neuroticism general factor. This modest intersection suggests that our inferences of enrichment by these two collections of SNPs were mostly independent.

The similarity of the biology implicated by general-factor and independent-pathway SNPs has two possible interpretations. First, the general factor and non-factor influences on the questionnaire items may tend to act through similar biological mechanisms. Second, as suggested by the concordance of effect signs observed in the GWAS of the group factors, it may be that the general factor is in fact one of several mechanisms affected by an independent-pathway SNP, the other mechanisms being responsible for the departures from the strict predictions of the general-factor model (Fig. 1A). To investigate the latter possibility, we calculated sign concordances of the SNP effects on the 12 items. Of the 181 SNPs, 117 showed sign-concordant effects on all 12 items, 28 showed a deviant sign with respect to only one item, 15 showed deviant signs with respect to two items, 11 showed deviant signs with respect to three items, and 10 showed deviant signs with respect to four items. The overall impression is that many of these SNPs do not depart too radically from the general-factor model, despite a low Akaike weight for the precise predictions of that model.

The Supplementary Data contain information about all of the SNPs used in these analyses.

### 3.6 S-LDSC and PANTHER fold enrichment

The apparent rarity of severe model failures among the more significant SNPs associated with the neuroticism general factor lends interpretability to genome-wide estimates of heritability enrichment, as calculated by S-LDSC, where there has been no screening of SNPs for conformity to the general-factor model (Fig. 1A).

It is recommended that S-LDSC be used with a standard collection of control variables. The estimates associated with these variables can be interesting in their own right, and we give them in Supplementary Table S14. The most statistically significant enrichments were shown by annotations referring to evolutionary conservation, more recent mutational origin, and lower correlations with nearby SNPs. This pattern is typical of traits that have been studied in GWAS (Finucane et al., 2015; Gazal et al., 2017). What the pattern means is that mutations affecting the neuroticism general factor (and other traits) tend to arise in functional regions of the genome, as evidenced by selection to maintain sequence similarity in distinct lineages, and once arisen tend to be deleterious.

Fig. 3 displays the enrichment estimates for the reconstituted gene sets and tissues/cell types. We first discuss the colored data points representing the tissues/cell types. The top result by heritability enrichment, as by DEPICT *p* value, was *entorhinal cortex* (1.45-fold enrichment, *p* = 10*^−^*^15^). Furthermore, 11 of the 13 tissues/cell types clearing 1.3-fold enrichment bore the MeSH second-level term *central nervous system*. The two non-CNS tissues/cell types were *neural stem cells* (1.31-fold enrichment, *p* = 1.9 *×* 10*^−^*^6^) and *retina* (1.31-fold enrichment, *p* = 6.4 *×* 10*^−^*^6^). These are not true exceptions. Neural progenitors are reasonably classified as neural despite differences in gene expression between progenitors and differentiated cells, and the retina is made up of layers of neurons.

**Figure 3:**
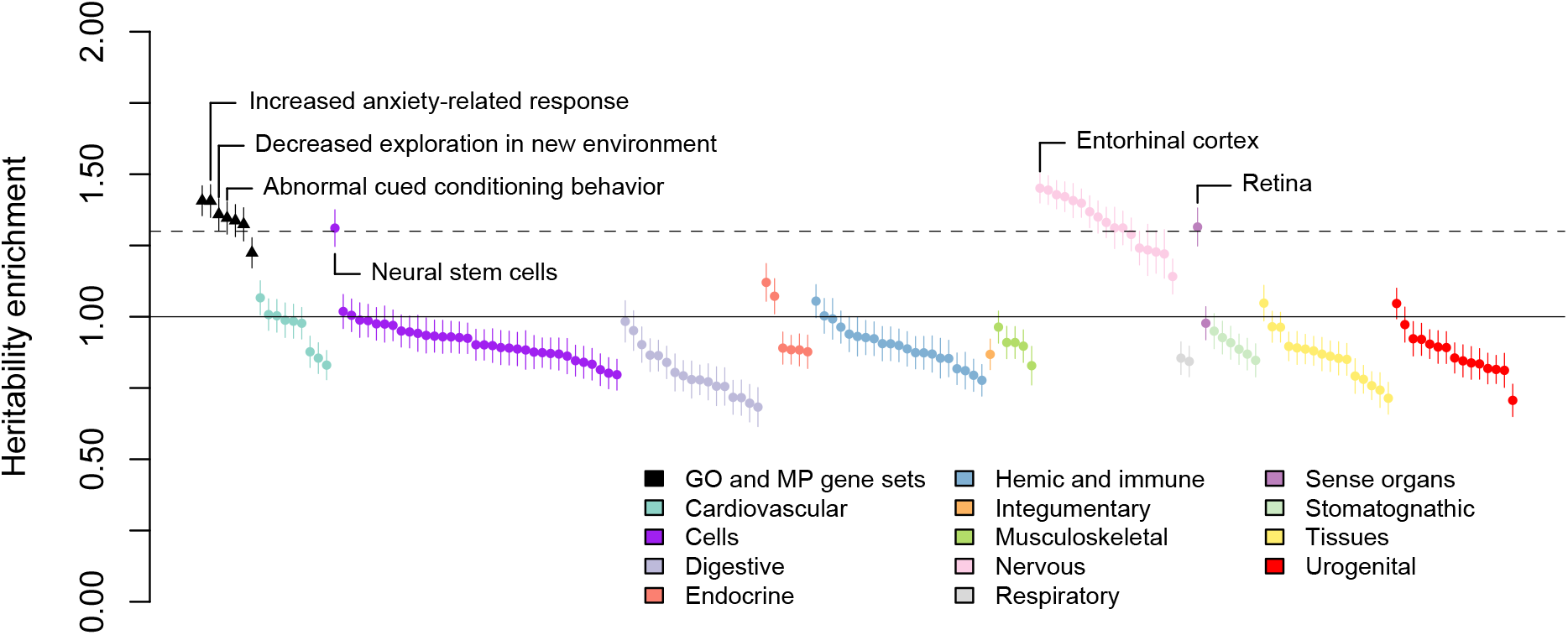
Heritability enrichment of reconstituted gene sets and tissues/cell types, as estimated by stratified LD Score regression (S-LDSC) applied to the GWAS summary statistics of the neuroticism general factor. The error bars are *±*1-SE intervals. The height of the dashed horizontal line corresponds to 1.3-fold enrichment, which we consider to be a “large” effect size. Complete numerical results are given in Supplementary Table S15. GO, Gene Ontology; MP, Mammalian Phenotype.

Before proceeding further, we point out that the any-gene control annotation typically showed roughly 1.03-fold enrichment with a standard error of .015, demonstrating that the S-LDSC estimation procedure is well calibrated.

We now turn to the reconstituted gene sets, which are represented by the dark data points at the far left of Fig. 3. All but *abnormal neuron physiology* (1.22-fold enrichment, *p* = 3.3*×*10*^−^*^5^) exceeded the benchmark effect size of 1.3. In particular, the gene sets defined in one way or another by fearful and anxious behavior in mice all met the threshold: *increased anxiety-related response* (1.41-fold enrichment, *p* = 4.1 *×* 10*^−^*^11^), *decreased exploration in new environment* (1.36-fold enrichment, *p* = 10*^−^*^9^), and *abnormal cued conditioning behavior* (1.35-fold enrichment, *p* = 3 *×* 10*^−^*^9^).

The PANTHER overrepresentation test also supported the results of the standard DEPICT analysis. Supplementary Table S16 shows that both *growth cone* (14.5-fold enrichment, *p* = 2.7 *×* 10*^−^*^5^) and *site of polarized growth* (14.1-fold enrichment, *p* = 3.1 *×* 10*^−^*^5^), the two gene sets in Table 2 related to axonogenesis, were among the top Gene Ontology (GO) cellular components. The theme of axonogenesis was reinforced by many of the significant GO biological processes (*neuron projection morphogenesis*, *axonogenesis*, *axon development*).

Because our pipeline of DEPICT-prioritized genes to PANTHER did not require defining a single effect size for any given SNP, we were able to perform the PANTHER overrepresentation test on prioritized genes near independent-pathway SNPs. Supplementary Table S17 shows that many gene sets reaching statistical significance for the neuroticism general factor also did so for independent pathways (e.g., *axon guidance*, *axon development*, *neuron projection*). Many of the most strongly enriched gene sets for independent pathways were defined by synaptic function: e.g., *presynapse assembly* (70.9-fold enrichment, *p* = 9.7 *×* 10*^−^*^10^), *synaptic vesicle clustering* (82.7-fold enrichment, *p* = 4.2 *×* 10*^−^*^7^), *neuron to neuron synapse* (7.6-fold enrichment, *p* = 6.9 *×* 10*^−^*^8^), *postsynaptic density* (7.2-fold enrichment, *p* = 1.6 *×* 10*^−^*^6^), *GABA-ergic synapse* (13.9-fold enrichment, *p* = 3.8 *×* 10*^−^*^5^) and *glutamatergic synapse* (5.2-fold enrichment, *p* = 6.8 *×* 10*^−^*^5^).

## 4 Discussion

The common-factor model need not be interpreted as a causal account of the correlations between indicators in order to be scientifically and practically useful (Ashton & Lee, 2005; McDonald, 1996, 2003). Nevertheless the extent to which factors do approximate underlying causes is a matter worthy of investigation.

Our results suggest that the factor model of the neuroticism domain is not just a convenient summary of the correlations between items, but indeed a reasonable approximation to some part of the underlying causal system. For instance, neuroticism does not appear to be explained entirely by something like the bonds model (Thomson, 1951), which proposes the existence of many distinct causal elements, no single one of which affects all items in the domain. In Thomson’s model, items may overlap in what bonds affect them, and a greater overlap produces a greater correlation. A resulting positive correlation between each pair of items then gives the appearance of a single causal variable affecting all items when in fact there is no such variable. Bartholomew et al. (2009) suggested that polymorphic sites in the human genome might turn out to be the substantiation of the abstract bonds in Thomson’s model, but our results show that many SNPs identified in a GWAS of a neuroticism questionnaire are in fact associated with all items as if mediated by the common factors.

Even upon rejecting a simpler model of mediation, we still found evidence for the approximate correctness of such a model. SNPs ascertained through a GWAS of the three group factors were found to show sign-concordant effects on those factors. In summary, we have genetic evidence supporting the verisimilitude of the neuroticism general factor at a deep biological level. This evidence weighs against network theories that deny the existence of broad factors influencing many specific traits (Cramer et al., 2012), adding specific neurobiological reasons to other statistical and theoretical reasons to reject such models as sufficient explanations of personality structure (DeYoung & Krueger, 2018).

We concede that our study cannot be absolutely definitive on this point. The filtering of SNPs by statistical significance in a GWAS at the latent level may induce an ascertainment bias that exaggerates the evidence for the approximate validity of the factor model. That is, SNPs departing very markedly from concordance of associations with all of the questionnaire items may be less likely to reach the threshold of statistical significance in a GWAS of the common factor. An example might be a SNP with positive effects on half of the items and negative effects on the other half. Such a SNP might have no net effect on the sum score and presumably would not reach significance in a GWAS of the factor, but it would be detected in a sufficiently powerful GWAS of independent pathways. Future research may attend to this issue of ascertainment bias more carefully. Again, however, it is telling that most of the SNPs ascertained solely for significant association with just one group factor showed evidence of concordant association with the two others as well. Regardless of what we have failed to ascertain, it is clear that there are a sizable number of polymorphic sites across the genome that bear a striking resemblance to causes of the neuroticism general factor.

A true GWAS of independent pathways, testing all common SNPs rather than those first attaining significance in a GWAS of a common factor, is likely to identify roughly as many lead SNPs as a GWAS of the general factor carried out on the same item-level summary statistics. For example, in their specification, Grotzinger et al. (2019) identified 118 lead SNPs for the neuroticism factor and 69 lead SNPs for independent pathways. Such a GWAS is also likely to identify more lead SNPs showing a failure of sign concordance across items. A tally of lead SNPs, however, may not suffice to weigh the relative importance of mechanisms. For example, if many independent-pathway lead SNPs are associated with item-specific residuals, then such SNPs are not in fact contributing to the correlational structure of this personality domain. We do not propose a suitable comparative metric at the current time, leaving this problem to be addressed in future research.

Previous studies have used multivariate twin modeling to pursue aims similar to our own. For example, Heath et al. (1989) showed that data from 1,800 pairs of like-sex monozygotic twins and 1,103 like-sex dizygotic twins were consistent with some personality scales being influenced by a general heritable factor. In their study this was true of extraversion and neuroticism, but not the third EPQ trait of psychoticism. This work may have contributed to the decline in support for the construct validity of psychoticism, showing the potential impact of genetic methods on personality theory. Even the fit of genetic correlations to a single factor, however, does not rule out a network or Thomson-like model. The power of the genomic approach lies in subjecting a factor model to an even more precise and hence riskier quantitative test of how directly measurable objects are related to the trait indicators (Meehl, 1978).

We applied DEPICT in order to gain some clues to the biological processes mediating the effects of the general-factor SNPs on neuroticism. We found that these SNPs disproportionately fall within or near genes designated as high-ranking members of gene sets defined by responses to aversive or novel stimuli (Table 2). This result is remarkably fitting for the personality trait of neuroticism. Such gene sets became significantly enriched in GWAS of other behavioral traits as their sample sizes grew (e.g., Lee et al., 2018b), but it is perhaps meaningful that they are among the first to become significantly enriched in the GWAS of a trait defined by a tendency to experience fear and anxiety. Furthermore, the tendency of these genes to be highly expressed in the entorhinal cortex (Fig. 2) is consistent with research and theory linking anxiety and the mechanisms of anxiolytic drugs to the septo-hippocampal system (Allen & DeYoung, 2017; Gray & McNaughton, 2000)—a collection of structures that receive from the medial septal nucleus the inhibitory GABAergic input inducing the theta rhythm, a neural oscillation associated with learning and spatiotemporal encoding in many animals. The main components of the septo-hippocampal system are the hippocampus itself and the entorhinal cortex (Robinson et al., 2023). The Cybernetic Big Five Theory (DeYoung, 2015), drawing on Gray and McNaughton (2000), posits that neuroticism reflects the joint sensitivity of a behavioral inhibition system (BIS), which responds to threats in the form of conflicts between goals (e.g., approach-avoidance conflict or any conflict that generates uncertainty), and a fight-flight-freeze system (FFFS), which responds to threats without conflict—that is, when the only motivation is to escape or eliminate the threat. Much is known about the neurobiology of the BIS and FFFS in the brainstem, hypothalamus, and limbic system [a collection of structures including the hippocampus and entorhinal cortex], which can aid in the interpretation of existing research on [n]euroticism and inform hypotheses in future research. (Allen & DeYoung, 2017, p. 331)

By and large, our biological-annotation results were consistent with previous analyses. For example, they were broadly consistent with those obtained with a different software tool, MAGMA (de Leeuw et al., 2015), in a GWAS of the questionnaire sum score (Nagel et al., 2018a). The three independently significant gene sets in this study were *neurogenesis*, *behavioral response to cocaine*, and *axon part*. Biological annotation apparently tends to yield similar results regardless of whether it is applied to the general factor, the observed sum score, or a single factor in a simpler model. Perhaps such consistency is to be expected in light of our evidence for the existence, in some sense other than the psychometric one, of a general factor. A sum score will typically reflect a general factor indicated by all items more than any other source of variance. Indeed, on the basis of the phenotypic correlations between items reported by Nagel et al. (2018b), we calculated McDonald’s *ω_H_* (Revelle & Condon, 2019) of the EPQ neuroticism scale to be .64.

We have no explanation for the meager results obtained from the GWAS of the residual group factors. Our method for identifying SNPs associated with the residuals of the group factors in our hierarchical model was somewhat indirect (Supplementary Fig. S2), but a more direct approach based on a bifactor model would lead to more free parameters and an increase in estimation error (Murray & Johnson, 2013; Preacher et al., 2013). The study of group factors is an inherently difficult one, and those present in the EPQ neuroticism questionnaire require a greater GWAS sample size for their genetic elucidation. It would be premature to base conclusions about the construct validity of these group factors on the present results.

## 5 Conclusion

We used structural equation modeling to carry out a GWAS of the neuroticism general factor and identified 19 lead SNPs satisfying *p <* 5 *×* 10*^−^*^8^. Even if deemed not to satisfy the predictions entailed by the hypothesis of acting solely through the general factor, hundreds of other SNPs attaining or approaching statistical significance in various analyses showed mostly sign-concordant effects on the questionnaire items. These findings do not settle the issue of the causal structure underlying the correlations between personality items. All we claim is that when we look for evidence of genetic effects on a causal intermediary very similar to the general factor of neuroticism, such evidence can be found. The SNPs acting through the general factor are found in or near genes highly expressed in the brain, and their pattern of gene-set enrichment is suggestive of neural development and synaptic function, particularly as these processes affect the learning of fear and caution in response to aversive stimuli.

## Data Availability

The Supplementary Data archive contains R code, several files containing limited portions of the Genomic SEM output, and a DEPICT configuration file. The original item-level GWAS summary statistics are available at https://ctg.cncr.nl/software/summary_statistics. The GWAS summary statistics generated for this paper and the DEPICT collections file used in our analyses are available at XXX.

