## supplementary material for "Genetic and neural bases of the neuroticism general factor"

**2 Departures from the analysis of Grotzinger et al. (2019)** **5**

**3 Differences between exploratory and confirmatory factor models** **6**

**4 Genetic correlations** **7**

**5 Polygenic prediction** **8**

**6 Remarks on cross-ancestry replicability** **9**

#### 1 Additional methods

##### 1.1 Original GWAS of neuroticism items

The original item-level GWAS were conducted by Nagel et al. (2018b), whose description of their methods is largely reproduced here.

The UK Biobank (UKB) is a major data resource, containing phenotypic measures from 503,325 participants and genetic data from 489,212 participants. The data were released in two phases (May 2015 and July 2017). Sample 1 consists of individuals for whom the data was released in May 2015 ($N = 110,328$), whereas sample 2 consists of all individuals that were added in July 2017 ($N = 270,178$). Written informed consent was obtained from all participants, and the UK Biobank received ethical approval from the National Research Ethics Service Committee North West—Haydock.

All individuals of non-European ancestry were excluded. In detail, UKB participants with genetic data were projected onto the principal components (PCs) from the 1000 Genomes reference populations. Those participants whose projected PC scores were closest to the average score of the Europeans in 1000 Genomes (based on the Mahalanobis distance) were identified as European. European subjects with a distance $> 6$ standard deviations were then excluded. Additionally, subjects were filtered out based on relatedness, discordant sex, sex aneuploidy, and withdrawn consent. Finally, individuals were excluded from analyses if they did not respond to 3 or more of the 12 items making up the neuroticism component of the Eysenck Personality Questionnaire–Revised Short Form (EPQ). The final sample size of the item-level GWAS was 380,506 individuals (205,556 females and 174,950 males). At the time of assessment, participants ranged in age between 40 and 73 ($M = 56.91$; $SD = 7.93$).

For both samples specified above, newly imputed data from the second release on $\sim 96$ million genetic variants was used. Imputation was performed by the UKB, using a reference panel that included the UK10K haplotype panel as well as the Haplotype Reference Consortium reference panel. Based on recommendations by the UKB, the authors excluded all variants imputed from the UK10K reference panel, as technical errors may have occurred during the imputation process. The imputed data were converted to hard-called genotypes using a certainty threshold of .9.

GWAS were conducted separately on samples 1 and 2 with logistic regression as implemented in PLINK 1.9 (Chang et al., 2015; Purcell et al., 2007). The dichotomous item responses were regressed on the imputed hard-called SNPs. Sex, age, and the Townsend Deprivation Index (measure based on postal code indicating material deprivation) were included in the analyses as covariates. Because of the genetic correlation between the sexes of .91 in the neuroticism sum score (Wendt et al., 2023), it was justified to analyze both sexes together. Genotype array was only included as covariate in the analyses of sample 1, as the same array was used for all subjects in sample 2. Additionally, the first 10 genetic PCs were used as covariates to control for potential population stratification. Genetic PCs were computed separately for both samples using FlashPCA 2 on individuals of European ancestry, after LD pruning and filtering out SNPs with MAF $< .01$, and genotype missingness $> .05$.

Final data analysis was restricted to autosomal, bi-allelic SNPs with MAF $> .0001$, high imputation quality (INFO score $\geq .9$) and low missingness ($< .05$).

#### 1.2 GWAS with Genomic SEM in the present study

The GWAS summary statistics produced by Nagel et al. (2018b) in the manner described above were given as input to LD Score regression (LDSC), as called by Genomic SEM, to calculate the genetic correlations between items. Standard procedures were followed (e.g., including only HapMap3 SNPs with a minor allele frequency greater than .01). The item prevalences were taken from Nagel et al. (2018b). The genetic covariance matrix thus obtained was used in a GWAS of the common factors depicted in Fig. 1. All defaults of Genomic SEM were employed, including diagonal weighted least squares as the estimation method, adjustment of the univariate GWAS standard errors by their univariate LDSC intercepts, and unit-loading identification. We used the reference file supplied by Genomic SEM to retain only SNPs with a minor allele frequency (MAF) exceeding .005 in the 1000 Genomes European populations. This left more than 7 million SNPs in the GWAS.

*Mood* was chosen as the item with a fixed loading of unity on the factor of depressed affect, *nervous* as the item with a fixed loading of unity on the factor of worry, and *guilt* as the item with a fixed loading of unity on the factor of vulnerability. Depressed affect was the first-order factor chosen as the indicator with a fixed loading of unity on the second-order neuroticism general factor. This identification strategy is important for understanding the scaling of SNP regression coefficients in Table 1, Supplementary Table S5, and Supplementary Table S9. For example, the two path coefficients each equaling unity from neuroticism to *mood* mean that any SNP with an effect of $\beta$ on neuroticism has the same effect $\beta$ on *mood*. At the same time, there are residual genetic effects on depressed affect and then on *mood*. Since the standardized loading of *mood* on depressed effect was estimated to be .966 (Supplementary Table S3), about 93 percent of the genetic variance in *mood* was attributed to depressed affect. In turn, since the standardized loading of depressed affect on neuroticism was estimated to be .796, about 63 percent of that .93 (i.e., 59 percent) was attributed to neuroticism. The liability of *mood* was estimated by LDSC to have a common-SNP heritability of .084 (Supplementary Table S3), meaning that the heritability of *mood* contributed by neuroticism must be $.084 \times .59 \approx .049$. Therefore, all common SNPs across the genome affecting the neuroticism general factor are constrained to satisfy the relation

$$.049 \approx \sum 2 \times \text{MAF} \times (1 - \text{MAF}) \times \beta^2.$$

This summed additive genetic variance over SNPs will equal the naive common-SNP heritability in a typical non-latent GWAS, meaning roughly .10 if the phenotype is years of education or .20 if it is IQ (Lee et al., 2018). This explanation of the scaling should dispel any impression that the regression coefficients given in our study (e.g., Table 1) are unusually small.

PLINK 1.9, as called by DEPICT, was used to identify roughly independent lead SNPs. The SNP with the lowest $p$ value in a given GWAS was chosen as the first lead SNP. All other SNPs less than 500 kb from this lead SNP and correlated with it to the extent $r^2 > .1$ was assigned to the SNP's "clump." The next clump was greedily formed around the SNP with the next lowest $p$ value not already assigned to the first clump. This process was iteratively continued until there were no more SNPs in the GWAS satisfying the recommended $p < 10^{-5}$. To construct a locus around a given lead SNP, the farthest SNP on a given side whose correlation with the lead SNP satisfied $r^2 > .5$ was designated the endpoint on that side. Overlapping loci according to this definition were merged.

#### 1.3 Genetic correlations with the residual group factors

The group factors in the hierarchical model have such high loadings on the general factor that any genetic correlations with the group factors are bound to resemble those with the general factor (Supplementary Table S3). We therefore attempted to use a bifactor model in order to approximate genetic correlations with the residuals of the group factors within the hierarchical model. Note that the bifactor is nested within the hierarchical model, the former being obtained from the latter by freeing implicit proportionality constraints on the factor loadings. At least in intelligence research, the fit of the bifactor model is often only somewhat better than the fit of the hierarchical model (e.g., Gignac, 2006).

We first tried a model including all three group factors: depressed affect, worry, and vulnerability. We encountered a number of difficulties, as often happens in attempting to fit the bifactor model. Fitting the model took about 15 times as long as fitting the corresponding hierarchical model, rendering it impractical for a GWAS over millions of SNPs. There were also a number of Heywood cases—indicators with estimated non-positive residual variances. We tried to alleviate the difficulties by dropping the group factor of vulnerability. Since in the corresponding hierarchical model this factor was found to have little residual genetic variance, we anticipated that dropping the factor would degrade the fit negligibly while greatly speeding up convergence. This proved to be the case; the SRMR increased from .044 to .048, while the convergence time improved to the point of being competitive with that of the hierarchical model. Supplementary Fig. S3 displays our bifactor model in its final form. The item *nervous* was estimated to have a negative residual variance, a difficulty that we patched up by fixing this variance to zero.

We ran the GWAS based on the path model in Supplementary Fig. S3 to produce the summary statistics needed to calculate genetic correlations with the two remaining group factors. We observed that the mean $\chi^2$ declined considerably in moving from the hierarchical to the bifactor model; in the latter, the mean $\chi^2$ of the general-factor GWAS was only 1.50, no better than shown by some of the items (Supplementary Table S4). The genetic correlation between the general factors in these models was .93.

Despite the difficulties in fitting the bifactor model, we judged the results adequate for calculating genetic correlations of other traits with depressed affect and worry.

#### 1.4 Polygenic prediction

To convert our GWAS summary statistics into weights for polygenic scores (PGS), we used the software tool PRS-CS (Ge et al., 2019). This tool converts the univariate regression coefficients obtained in the GWAS into partial regression coefficients by plugging in a SNP covariance matrix taken from a reference panel and applying Bayesian continuous shrinkage rather specifying a discrete number of prior distributions. We used the 1000 Genomes Project phase 3 participants of European descent as the reference panel.

Our validation sample consisted of the Minnesota Twin Family Study and the Sibling Interaction and Behavior Study, both of which are being conducted by the Minnesota Center for Twin and Family Research (MCTFR) (Wilson et al., 2019). Each complete unit in the validation sample was made up of two siblings (usually twins) and their parents. The sample consisted of 9,067 total individuals belonging to 2,497 family units. Miller et al. (2012) provided details about genotyping and quality control.

The validation sample was not given a standard measure of Big Five neuroticism. The closest such measure that was in fact administered to the sample was the "negative emotionality" component (excluding the Aggression primary scale) of the Multidimensional Personality Questionnaire (MPQ) (Church, 1994; Tellegen & Waller, 2008). Item response theory was employed to estimate a factor score for each participant (van den Berg et al., 2014).

To benchmark the predictive power of our PGS, we compared it to a distinct PGS previously computed by Becker et al. (2021) for use in the same validation sample. These authors drew upon a GWAS of the neuroticism sum score performed by Nagel et al. (2018a) for their summary statistics and used the software tool LDpred (Vilhjálmsson et al., 2015) to calculate the PGS weights. PRS-CS and LDpred are based on similar principles and should perform about equally well in our application.

Ten genetic PCs, sex, age, and the square of age were used as covariates when regressing negative emotionality on the PGS. To deal with dependence between siblings in the same family, we performed bootstrap resampling over families to calculate standard errors. A thousand bootstrap replicates were used.

We repeated all of our PGS predictions except restricting observations to individuals with genotyped parents and adding the parental PGS as covariates. For a fixed value of the parental PGS, the PGS of the offspring vary randomly as a result of Mendelian segregation and thus provide a strong degree of causal inference (Laird & Lange, 2006; Lee, 2012; Lee & Chow, 2013; Okbay et al., 2022). There were only 2,056 members of the offspring generation in our sample with two genotyped parents.

#### 2 Departures from the analysis of Grotzinger et al. (2019)

Our work extends Grotzinger et al. (2019), in a manner that we now explain.

Supplementary Figure 4 of Grotzinger et al. shows what was done in that paper. The authors performed a GWAS specifying a single neuroticism factor measured by all items. They also performed an independent-pathways GWAS and identified 69 SNPs fitting the independent-pathways model better than one where the SNP acts through the single factor, at the significance threshold $p < 5 \times 10^{-8}$. They then examined whether these 69 SNPs would continue to fit the independent-pathways model better if the more parsimonious model was one where the SNP acts through two or three factors.

The authors found that for each additional factor posited in the model, there was a reduction in the number of SNPs showing a significantly better fit to the independent-pathways model. This pattern by itself strongly suggests that a model of a SNP acting through common factors rather than independent pathways will tend to fit better as the fit of the factor model itself improves. Note that the SRMR dropped from .109 to .057 as the number of factors in the model went from one to three. In our view an SRMR exceeding .10 is indicative of a poor fit, which we confirmed by finding several large elements in the residual correlation matrix resulting from a one-factor model.

All that said, the Grotzinger et al. strategy of beginning with a one-factor model and then testing SNPs deviating from that model within more accurate factor models may not be an unreasonable way to deal with the bias-variance tradeoff that penalizes greater model accuracy with a loss of GWAS signal (Supplementary Table S4). One might worry, however, that a simple model with poor fit might lead to an excess of independent-pathway SNPs with growing GWAS sample size. Whatever the strategy adopted, it is clear that any attempt to pit common- and independent-pathway models against each other must take into account the multidimensional basis of the factor space at some point in the procedure.

The construct validity of a general factor seems not to have been a main concern of Grotzinger et al. They did not specify a general factor in addition to the group factors in their two- and three-factor models. The authors mentioned performing a GWAS over HapMap3 SNPs of the two correlated factors and of the three correlated factors, but to our knowledge have not detailed or deposited these results anywhere. They did not perform biological annotation of their multiple-factor results. Even their biological annotation of their one-factor results was somewhat limited because they only provided $p < 5 \times 10^{-8}$ lead SNPs as input to DEPICT, whereas the developers of this tool recommend a more liberal threshold of $p < 10^{-5}$. As a result Grotzinger et al. found only one gene set to be significantly enriched.

In summary, we extended a model of three correlated factors by converting it to a hierarchical model with a second-order general factor and followed up a GWAS based on this model with the bioinformatic tool DEPICT. The latter tool was set to the developer-recommended parameter values. We also followed up the DEPICT results with additional bioinformatic analyses yielding effect sizes in terms of fold enrichment.

### 3 Differences between exploratory and confirmatory factor models

A reviewer expressed concern over differences between the results of the exploratory factor analysis reported in Supplementary Table S2 of Grotzinger et al. (2019) and those of the confirmatory model reported in their Supplementary Figure 3. The reviewer pointed, as an example, to the large cross-loading of the item *guilt* on an additional factor only in the exploratory analysis. Note that our confirmatory hierarchical model is equivalent to the confirmatory Grotzinger et al. (2019) model of three correlated factors.

Any such differences need not be an indication of serious problems, because what often occurs if one fits a confirmatory factor model that has been inspired by an exploratory analysis is that the correlations between factors increase as a way of compensating for the loss of fit induced by fixing many loadings to zero in order to form an independent-clusters solution (sometimes called "simple structure") (McDonald, 1999). This is exactly what happened in the application of Grotzinger et al. (2019). In the exploratory solution the correlations between factors were .42, .57, and .62, whereas in the confirmatory solution they were .73, .67, and .84. The fit of the confirmatory model was good (see the fit indices reported in the main text), testifying to the "success" of the tradeoff in fit.

We calculated the correlation matrix implied by the estimated parameters of the exploratory factor analysis in Supplementary Table S2 of Grotzinger et al. (2019) and compared it to the observed genetic correlation matrix. The SRMR was .044, which was only somewhat smaller than the .054 attained by our confirmatory hierarchical model. We went on to inspect the residual covariance matrix associated with our confirmatory model and did not observe any pattern of large residuals between *guilt* and other items. In short, we found no sign of any important features revealed by the exploratory analysis but left out of the confirmatory analysis.

#### 4 Genetic correlations

We calculated genetic correlations between the factors present in the EPQ neuroticism questionnaire and the following traits:

- subjective well-being (Okbay et al., 2016)
- household income (Hill et al., 2019)
- risk tolerance (Linnér et al., 2019)
- morning person (Hu et al., 2016)
- years of education (Lee et al., 2018)
- IQ (i.e., cognitive performance) (Lee et al., 2018)
- brain volume (Jansen et al., 2020)
- autism spectrum disorder (ASD) (Grove et al., 2019)
- schizophrenia (Pardiñas et al., 2018)
- bipolar disorder (Mullins et al., 2021)

- attention deficit/hyperactivity disorder (ADHD) (Demontis et al., 2019)
- obsessive compulsive disorder (OCD) (International Obsessive Compulsive Disorder Foundation Genetics Collaborative & OCD Collaborative Genetics Association Studies, 2018)
- major depressive disorder (Howard et al., 2019)
- number of children (Barban et al., 2016)
- number of sex partners (Linnér et al., 2019)
- drinks per week (Linnér et al., 2019)
- cannabis use disorder (CUD) (Johnson et al., 2020)

The pattern of results for the neuroticism general factor displayed in Supplementary Fig. S4—most notably the genetic correlations of large magnitude with subjective well-being and major depression—was indeed very similar to what has been obtained with observed measures of neuroticism (Baselmans et al., 2019; Luciano et al., 2018; Nagel et al., 2018a; Okbay et al., 2016; Turley et al., 2018).

Also of interest are the genetic correlations with the residuals of depressed affect and worry. The latter trait corresponds roughly to a factor called anxiety/tension by Hill et al. (2020). These authors found an intriguing tendency of neuroticism and anxiety/tension (i.e., what we are calling worry) to show genetic correlations of opposite sign with various traits in the domains of abilities, socioeconomic status, and health. We checked on whether this pattern was replicated in our study and found this to be roughly the case (Supplementary Fig. S4). For example, ADHD showed a positive genetic correlation with the neuroticism general factor ($r_g = .26$, $p < 10^{-8}$) but a negative genetic correlation with the worry residual factor ($r_g = -.19$, $p < .005$). Sometimes one genetic correlation was attenuated in magnitude without reversing sign. Note that the similarity between results cannot be considered a replication in the fullest sense because they are based on the same neuroticism data (i.e., the UKB), but it is notable that substantial methodological differences between studies did not obscure the basic pattern.

### 5 Polygenic prediction

Supplementary Table S2 presents the results. The PGS of Becker et al. (2021) based on the observed sum score was a highly significant predictor of negative emotionality ($p < .001$), but the incremental prediction $R^2$ was only .006, substantially smaller than what others have observed in similar exercises (e.g., Luciano et al., 2018). We do not have an explanation for this disparity, although one contributor might be an imperfect genetic correlation between the EPQ neuroticism questionnaire and MPQ negative emotionality.

Our PGS based on the neuroticism general factor outperformed the Becker et al. (2021) PGS, reaching an incremental prediction $R^2$ of .009 ($p < .001$). Despite the GWAS at the latent level showing a lower mean $\chi^2$ over HapMap3 SNPs (Supplementary Table S4), a PGS based on this GWAS might better capture the heritable influences on MPQ negative emotionality.

We added the PGS based on the group factors of depressed affect and worry to the regression model. We obtained little, if any, increment to the prediction $R^2$.

Because of the drop in statistical power, we could not draw many firm conclusions from the within-family prediction of MPQ emotionality. The regression coefficients of the PGS were plausibly the same at the individual level and within families. The PGS based on the neuroticism general factor reached marginal significance ($p < .05$).

### 6 Remarks on cross-ancestry replicability

The GWAS were conducted in European-ancestry individuals exclusively, as explained in a previous section. We expect that the results have roughly the same applicability to other ancestries as has been observed in other GWAS: strong concordance of the lead SNPs, especially those with higher minor allele frequencies, but a decline in the correlation of the polygenic score with the phenotype that is linear with the $F_{ST}$ of the two populations (Marigorta & Navarro, 2013; Scutari et al., 2016). Note that this decline can be very plausibly attributed in most cases to differences in allele frequencies and correlations between SNPs, without invoking a true moderation of the causal effects (Hou et al., 2023).

One exception to the general trend of GWAS replicability across populations might be major depression. We know of one GWAS of depression including both European and African Americans (Levey et al., 2021). This showed reasonable-seeming concordance between the two groups (see this paper's Figure 5b), but the sample size of the African Americans was too small to draw a strong conclusion. Giannakopoulou et al. (2021) conducted a GWAS of depression in East Asians and compared the results to those of a European GWAS. These authors did find a low genetic correlation between Europeans and East Asians of .41, albeit with a large standard error. In this case the difference in ancestry was confounded with a difference in region and possibly diagnostic practices.

The depression results tentatively suggest that the genetically correlated trait of neuroticism might possibly break from the general trend for GWAS results to transfer well across populations and provide a motivation to conduct GWAS of personality traits in different settings.

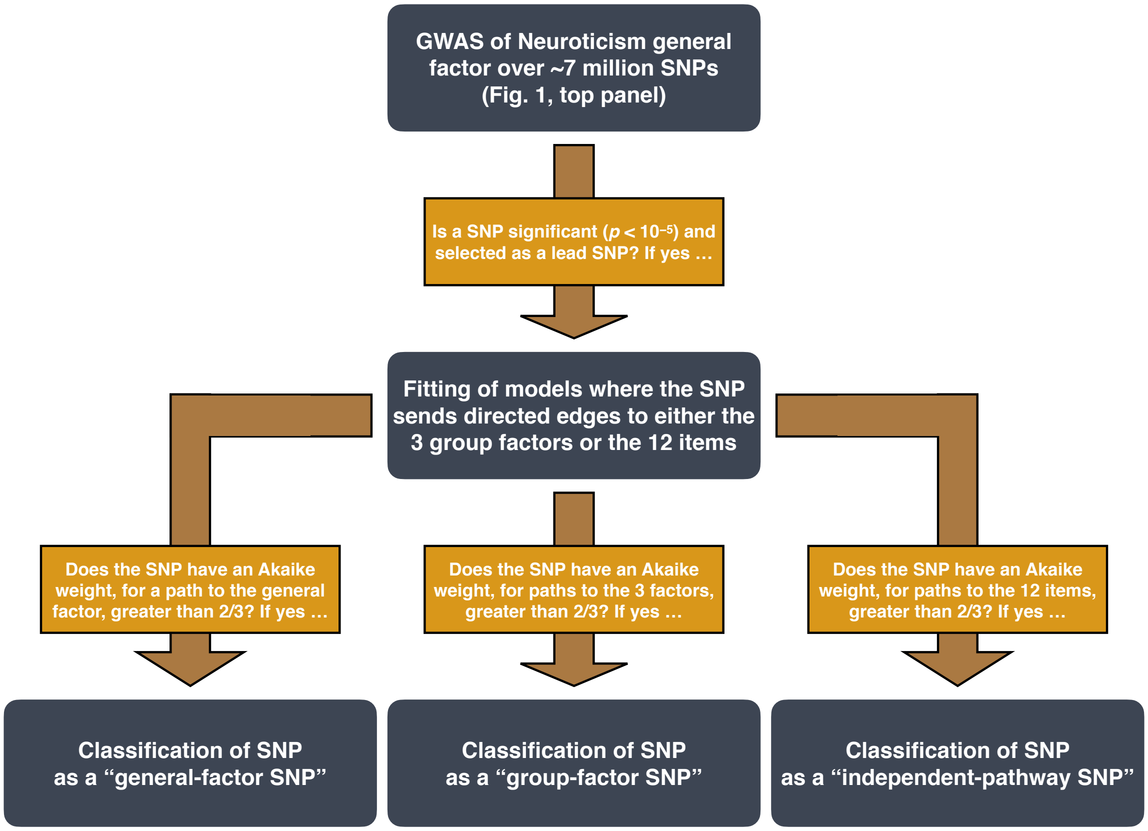

Figure S1: Flowchart of pipeline for our genome-wide association study (GWAS) of the Neuroticism general factor and subsequent classification of lead SNPs. Each gray box corresponds to an analysis step. Later steps use the output of earlier steps as input; such a relationship is represented by a gold arrow. The label of an arrow describes how the output of the prior step was filtered. Path modeling in the context of genetic association testing was conducted with Genomic SEM. Fig. 1 of the main text depicts the general-factor and independent-pathway model. Note that a SNP may not qualify for any of the three classifications in the final step.

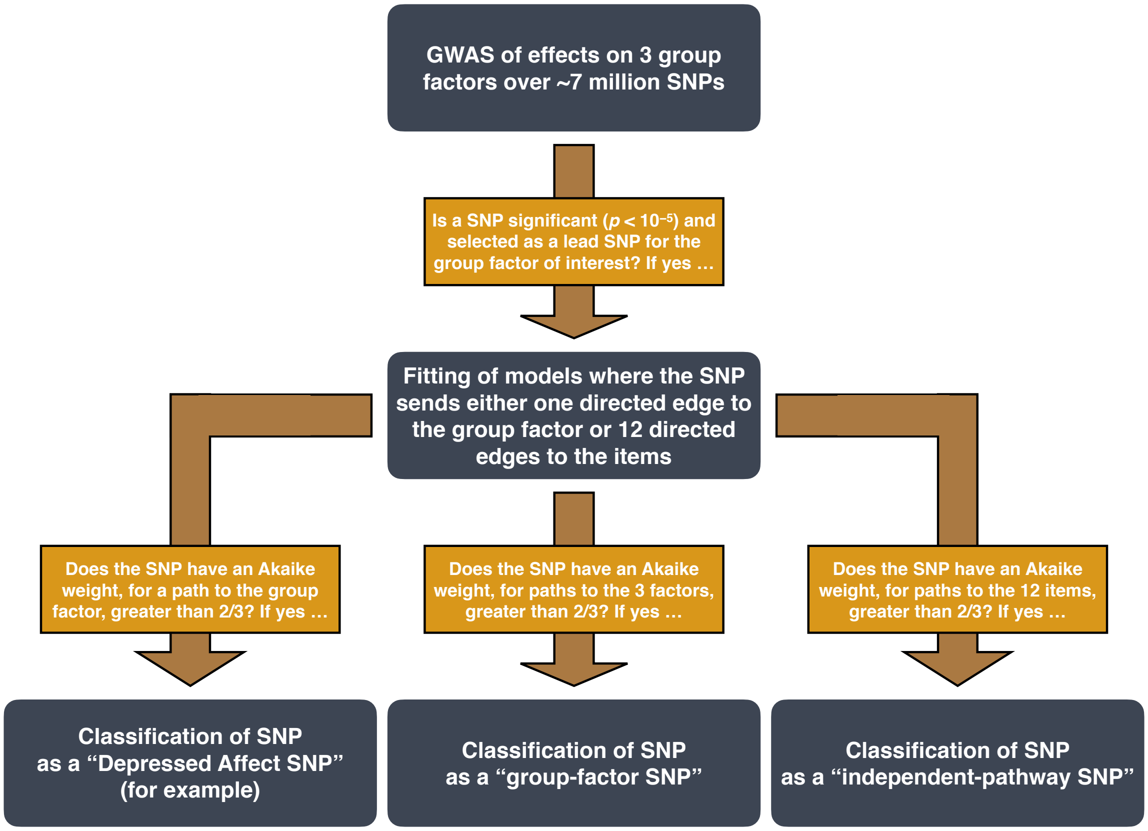

Figure S2: Flowchart of pipeline for our genome-wide association study (GWAS) of the group factors indicated by the Neuroticism items in the Eysenck Personality Questionnaire–Revised Short Form and our subsequent classification of lead SNPs. Each gray box corresponds to an analysis step. Later steps use the output of earlier steps as input; such a relationship is represented by a gold arrow. The label of an arrow describes how the output of the prior step was filtered. Path modeling in the context of genetic association testing was conducted with Genomic SEM. Fig. 1B of the main text depicts the independent-pathway model. Note that a SNP may not qualify for any of the three classifications in the final step.

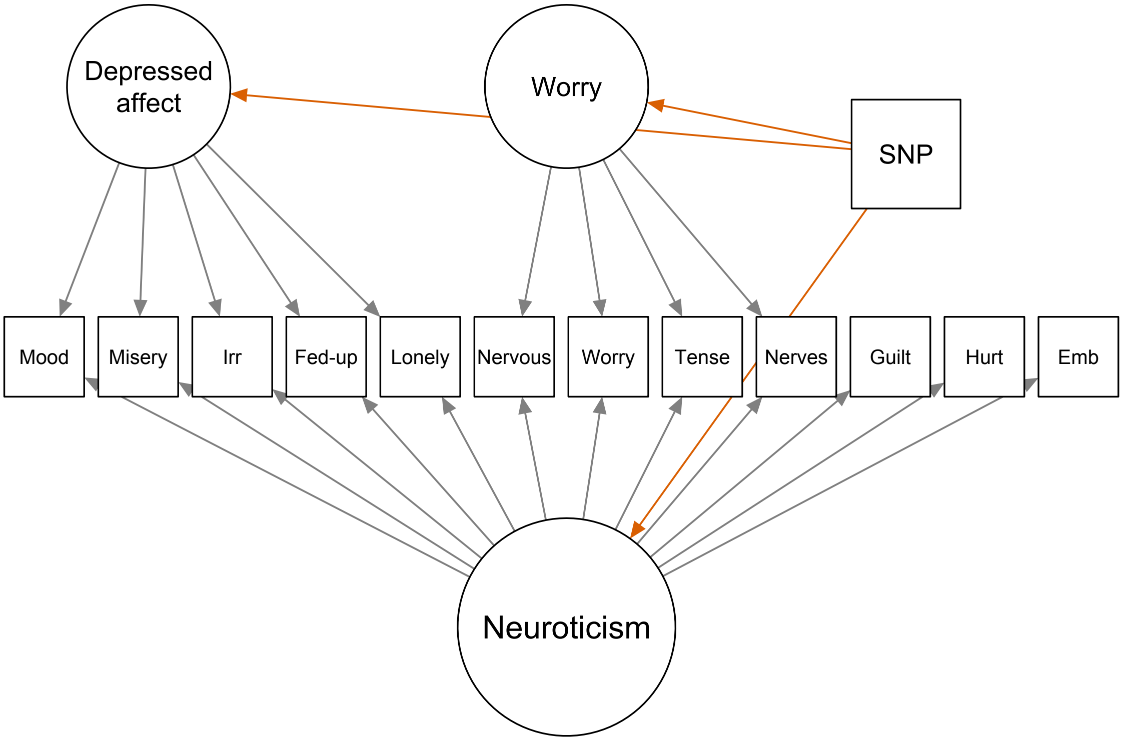

Figure S3: The bifactor model used to estimate genetic correlations with the group factors depressed affect and worry.

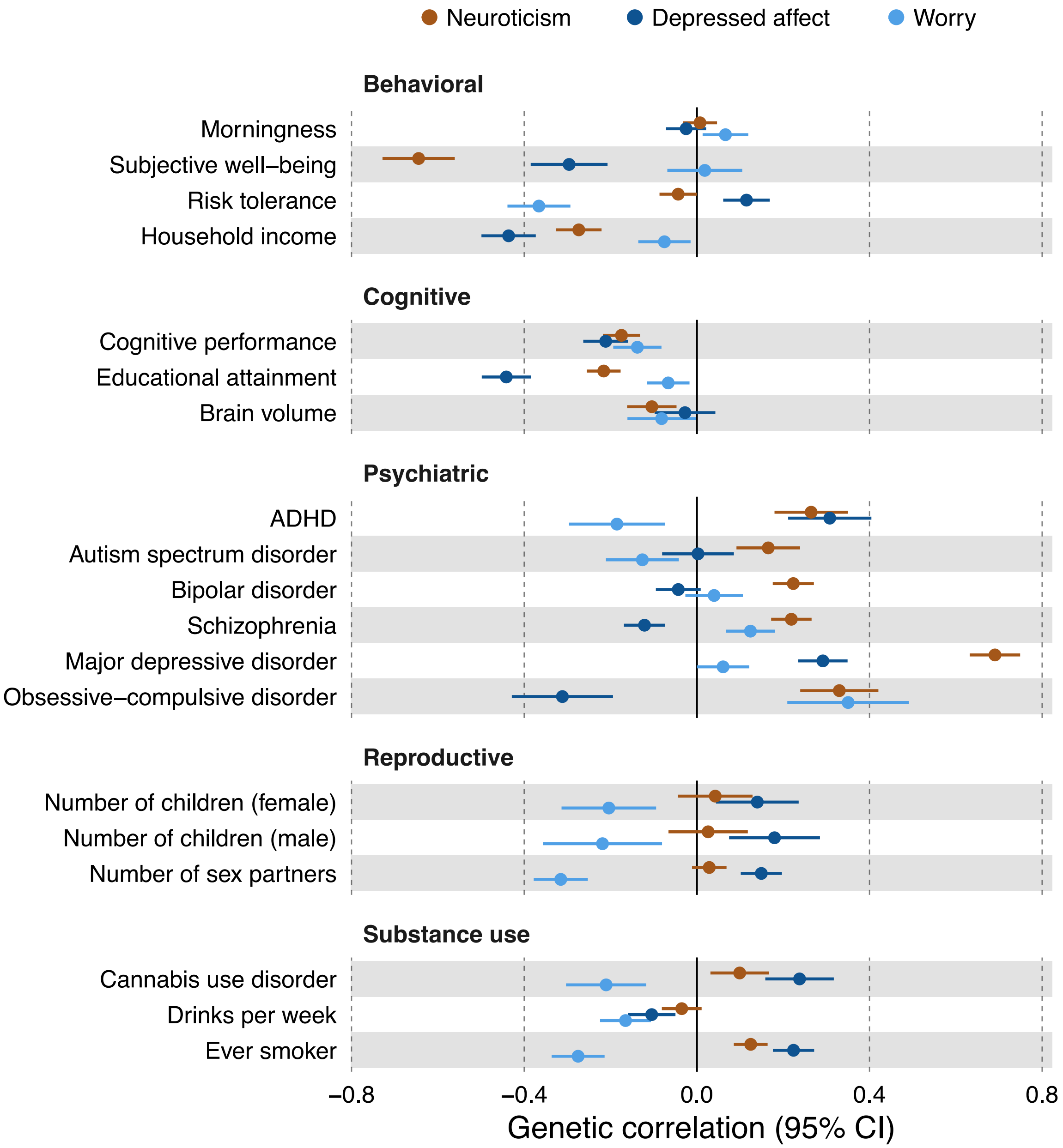

Figure S4: Genetic correlations with the neuroticism general factor and the residual group factors of depressed affect and worry. The estimates and accompanying ±1.96-SE intervals were calculated with LD Score regression, as called by Genomic SEM. Supplementary Table S1 gives the results in numerical form.
