## Supplementary material for "Genetic and neural bases of the neuroticism general factor": DEPICT_config_file.cfg

```
[PATHS]

# Absolute path to the output files will be written
analysis_path: # THE DIRECTORY WHERE YOU WANT THE DEPICT OUTPUT TO BE PLACED


[GWAS FILE SETTINGS]

# Absolute path to GWAS summary statistics (can be in plain text or gzip format) 
# gwas_summary_statistics_file: Neuroticism_model_3fact_GWASofN_allchr.txt.gz
# Use the summary statistics appropriate for your purpose. Indentifying lead SNPs,
# performing the bioinformatic analyses of a certain collection of SNPs, etc.
gwas_summary_statistics_file: N_lead_SNPs_AIC1.txt
# DO NOT FORGET TO PRECEDE THE FILE NAME WITH THE ABSOLUTE PATH

# Association cutoff used by DEPICT to define associated loci
association_pvalue_cutoff: 1e-5

# Label for output files generated by DEPICT
label_for_output_files: Neuroticism_model_3fact_GWASofN_AIC1_1E-5

# Name of p value column
pvalue_col_name: Pval_Estimate

# Name of marker name column. Format: <chr:pos>, ie. '6:2321'.  If this column does not exist chr_col and pos_col will be used, then leave if empty.
marker_col_name:

# Name of chromosome column. Leave empty if this column does not exist.
chr_col_name: CHR

# Name of chromosome column. Leave if this column does not exist.
pos_col_name: BP

# Separator options: 'tab', 'comma', 'semicolon' or 'space'
separator: tab


[PLINK SETTINGS]

# Change to your PLINK v. 1.9 August 5 release (or higher)
plink_executable: /Volumes/Backup_Plus/bioinformatics/plink_mac/plink

# Point this to the PLINK formated genotype data (change only the path to DEPICT if you use the 1000 Genomes data provided with DEPICT)
#genotype_data_plink_prefix: data/genotype_data_plink/CEU_GBR_TSI_unrelated.phase1_release_v3.20101123.snps_indels_svs.genotypes_ldl_teslovich_nature2010
genotype_data_plink_prefix: data/genotype_data_plink/CEU_GBR_TSI_unrelated.phase1_release_v3.20101123.snps_indels_svs.genotypes_noduplicates

# Options that typically will not be changed
plink_clumping_snp_column_header: SNP
association_pvalue_cutoff_column_header: P
plink_clumping_distance: 500 
plink_clumping_r2: 0.1
# These values used to be hardcoded into DEPICT. Now they can be changed. They must be specified.

[DEPICT SETTINGS]

# The following three steps are need to construct DEPICT loci based on your GWAS summary statistics
step_write_plink_input: yes
step_construct_depict_loci: yes

# The following step is needed to perform DEPICT gene prioritization
step_depict_geneprio: yes

# The following step is needed to perform DEPICT gene set enrichment
step_depict_gsea: yes

# The following step is needed to perform DEPICT tissue enrichment analysis
step_depict_tissueenrichment: yes


[MISC SETTINGS]

# Number of threads used by the DEPICT Java binary
number_of_threads: 12

# Java heap size in mega bytes
heap_size_in_mb: 16000

# Precomputed loci for each 1000 Genomes project SNPs
# collection_file: data/collections/ld0.5_collection_1000genomespilot_depict_150429.txt.gz
collection_file: data/collections/ld0.5_collection_1000genomesphase3_depict_170802.txt.gz

# The reconstituted gene set files used by DEPICT
reconstituted_genesets_file: data/reconstituted_genesets/reconstituted_genesets_150901.binary 
                                                        

# The tissue expression matrix
tissue_expression_file: data/tissue_expression/GPL570EnsemblGeneExpressionPerTissue_DEPICT20130820_z_withmeshheader.txt

# Mapping from tissue/cell type identifier to tissue name and information
tissue_mapping_file: data/tissue_expression/GPL570EnsemblGeneExpressionPerTissue_DEPICT20130820_z_withmeshheader_mapping.txt

# Gene annotation file
depict_gene_annotation_file: data/mapping_and_annotation_files/GPL570ProbeENSGInfo+HGNC_reformatted.txt

# Number of genes to report for each reconstituted gene set
max_top_genes_for_gene_set: 25

# Number of repititions used to compute false discovery rates
nr_repititions: 50

# Number of permutations used to adjust for biases such as gene length
nr_permutations: 500

# Boundaries of HLA region (Should not be changed)
mhc_start_bp: 25000000
mhc_end_bp: 35000000

# Directory with precomputed background files
background_data_path: data/backgrounds

# Mapping from gene set identifiers to gene set names
go_mapping_file: data/mapping_and_annotation_files/GO.terms_alt_ids_withoutheader.tab
mgi_mapping_file: data/mapping_and_annotation_files/VOC_MammalianPhenotype.rpt
inweb_mapping_file: data/mapping_and_annotation_files/inweb_mapping.tab

# Files used to output eQTL column in gene prioritization result file
eqtl_mapping_file: data/mapping_and_annotation_files/2012-08-08-IlluminaAll96PercentIdentity-ProbeAnnotation-ProbesWithWrongMappingLengthFilteredOut-EnsemblAnnotation.txt
eqtl_file: data/mapping_and_annotation_files/eQTLProbesFDR0.05.txt

# Prioritize genes across entire genome
prioritize_genes_outside_input_loci: yes

# Chromosome to be left-out, leave empty if all chromosomes should be included (for bencharmking purposes)
leave_out_chr:

# Number of null GWAS used to run permutations (bias adjustment) and repitions (FDR calcuations)
number_random_runs: 500

# Threshold used when clumping null GWAS
background_plink_clumping_pvalue: 0.0005
```
