## Supplementary material for "Genetic and neural bases of the neuroticism general factor": genomicSEM_revision_forpub.R

```
####################################################################################################

# install.packages("devtools")
# library(devtools)
# install_github("MichelNivard/GenomicSEM")

require(GenomicSEM)

###

GWAS_dir <- # THE LOCATION OF THE NEUROTICISM SUMMARY STATS AT THE ITEM LEVEL
data_dir <- # THE LOCATION WHERE YOU WILL PLACE OUTPUT FILES

item_names <- c(
"mood",
"misery",
"irrit",
"hurt",
"fedup",
"nervous",
"worry",
"tense",
"embarass",
"nerves",
"lonely",
"guilt")


# Run the GenomicSEM implementation of LDSC.

traits <- c(
paste(data_dir, item_names, ".sumstats.gz", sep="")
)

# These come from Supplementary Table 5 of Nagel et al. (2018)'s item-level paper.

sample.prev <- c(
0.45, # "mood",
0.43, # "misery",
0.28, # "irrit",
0.55, # "hurt",
0.40, # "fedup",
0.24, # "nervous",
0.57, # "worry",
0.18, # "tense",
0.48, # "embarass",
0.21, # "nerves",
0.18, # "lonely",
0.29 # "guilt"
)

population.prev <- c(
0.45, # "mood",
0.43, # "misery",
0.28, # "irrit",
0.55, # "hurt",
0.40, # "fedup",
0.24, # "nervous",
0.57, # "worry",
0.18, # "tense",
0.48, # "embarass",
0.21, # "nerves",
0.18, # "lonely",
0.29 # "guilt"
)

ldsc_files <- # LOCATION OF THE LDSC EUR_REF AND EUR_W LD_CHR FILES
ld <- paste(ldsc_files, "eur_ref_ld_chr/", sep="")
wld <- paste(ldsc_files, "eur_w_ld_chr/", sep="")
LDSC_output <- ldsc(traits, sample.prev, population.prev, ld, wld, item_names,
                    sep_weights=TRUE)
save(LDSC_output, file=paste(data_dir, "Neuroticism_items_LDSC_output.Rdata", sep=""))

load(paste(data_dir, "Neuroticism_items_LDSC_output.Rdata", sep=""))

model_3fact <-"F1 =~ mood + misery + irrit + fedup + lonely 
               F2 =~ nervous + worry + tense + nerves
               F3 =~ guilt + hurt + embarass
                N =~ F1 + F2 + F3"

model_3fact_out <- usermodel(LDSC_output, "DWLS", model_3fact, CFIcalc=TRUE, std.lv=TRUE)

# Reviewer 1 asks why not compare to a one-factor model.

model_1fact <- paste("N =~ mood + misery + irrit + fedup + lonely + ",
                     "nervous + worry + tense + nerves + ",
                     "guilt + hurt + embarass",
                     sep="")
model_1fact_out <- usermodel(LDSC_output, "DWLS", model_1fact, CFIcalc=TRUE, std.lv=TRUE)
# Fit is bad. SRMR almost doubles.

# Now to do a GWAS of all SNPs.

model_3fact_GWASofN <-"F1 =~ mood + misery + irrit + fedup + lonely 
                       F2 =~ nervous + worry + tense + nerves
                       F3 =~ guilt + hurt + embarass
                        N =~ F1 + F2 + F3
                        N ~ SNP"

item_ID <- seq(1920, 2030, 10)
filenames <- rep(NA, 12)
for(i in 1:12){
  filenames[i] <- paste(GWAS_dir, "sumstats_neuro_f", item_ID[i], "_ctg_format.txt.gz", sep="") }

neuroticism_sumstats <- sumstats(
  filenames, 
  paste(data_dir, "reference.1000G.maf.0.005.txt", sep=""),
  item_names,
  se.logit=rep(TRUE, 12),
  parallel=FALSE
)
save(neuroticism_sumstats, file=paste(data_dir, "Neuroticism_items_gSEM_sumstats.Rdata", sep=""))

####################################################################################################
### START ###

# Do not run the code in this block interactively. This will probably crash your GUI. Instead
# insert this block into a separate .R file and call it at the command line with Rscript.

require(GenomicSEM)

GWAS_dir <- # THE LOCATION OF THE NEUROTICISM SUMMARY STATS AT THE ITEM LEVEL
data_dir <- # THE LOCATION WHERE YOU WILL PLACE OUTPUT FILES
PLINK_call <- # THE LOCATION OF YOUR COPY OF PLINK 1.9

load(paste(data_dir, "Neuroticism_items_LDSC_output.Rdata", sep=""))
load(paste(data_dir, "Neuroticism_items_gSEM_sumstats.Rdata", sep=""))

model_3fact_GWASofN <-"F1 =~ mood + misery + irrit + fedup + lonely 
                       F2 =~ nervous + worry + tense + nerves
                       F3 =~ guilt + hurt + embarass
                        N =~ F1 + F2 + F3
                        N ~ SNP"

for(i in 1:22){
  print(i)
  GWAS <-  userGWAS(LDSC_output, subset(neuroticism_sumstats, CHR==i), "DWLS", model_3fact_GWASofN,
                    modelchi=FALSE, printwarn=FALSE, sub=c("N~SNP"), parallel=TRUE)
  filename <- paste(data_dir, "Neuroticism_model_3fact_GWASofN_chr", i, ".txt", sep="")
  write.table(GWAS, filename, sep="\t", row.names=FALSE, quote=FALSE)
  rm(GWAS)
  command <- paste("gzip ", filename, sep="")
  system(command) }

# WARNING
# The above loop might lead to exhaustion of memory, particularly after a larger chromosome.
# You might have to restart your machine if this happens. If you cannot execute this job on your
# local machine, please turn to whatever supercomputing resources are available to you.

setwd(data_dir)

# Insert the header into a new file that will contain the GWAS results from all chromosomes.
command <- paste("gzcat Neuroticism_model_3fact_GWASofN_chr22.txt.gz | ",
                 "head -n 1 > ",
                 "Neuroticism_model_3fact_GWASofN_allchr.txt",
                 sep="")
system(command)

for(i in 1:22){
  print(i)
  command <- paste("gzcat Neuroticism_model_3fact_GWASofN_chr", i, ".txt.gz | ",
                   "tail -n +2 >> ",
                   "Neuroticism_model_3fact_GWASofN_allchr.txt",
                   sep="")
  system(command) }

system("gzip Neuroticism_model_3fact_GWASofN_allchr.txt")

### END ###
####################################################################################################

# PLINK seems unable to clump the results in GenomicSEM format. We can try to get around this by
# converting the GWAS results to the format of a PLINK .assoc file. Another way is to let DEPICT
# do the clumping for us and get the results out of the .clumped or _loci.txt file.

####################################################################################################
### START ###

require(GenomicSEM)

GWAS_dir <- # THE LOCATION OF THE NEUROTICISM SUMMARY STATS AT THE ITEM LEVEL
data_dir <- # THE LOCATION WHERE YOU WILL PLACE OUTPUT FILES
PLINK_call <- # THE LOCATION OF YOUR COPY OF PLINK 1.9

load(paste(data_dir, "Neuroticism_items_LDSC_output.Rdata", sep=""))
load(paste(data_dir, "Neuroticism_items_gSEM_sumstats.Rdata", sep=""))

model_3fact_GWASofN <-"F1 =~ mood + misery + irrit + fedup + lonely 
                       F2 =~ nervous + worry + tense + nerves
                       F3 =~ guilt + hurt + embarass
                        N =~ F1 + F2 + F3
                        N ~ SNP"

model_3fact_GWASofF <- "F1 =~ mood + misery + irrit + fedup + lonely 
                        F2 =~ nervous + worry + tense + nerves
                        F3 =~ guilt + hurt + embarass
                        N =~ F1 + F2 + F3
                        F1 + F2 + F3 ~ SNP"

model_3fact_GWASindpath <- "F1 =~ mood + misery + irrit + fedup + lonely 
                        F2 =~ nervous + worry + tense + nerves
                        F3 =~ guilt + hurt + embarass
                        N =~ F1 + F2 + F3
mood + misery + irrit + fedup + lonely + nervous + worry + tense + nerves + guilt + hurt + embarass ~ SNP"


# The file .clumped contains lead SNPs in the GWAS of the Neuroticism general factor
# (model_3fact_GWASofN), as identified by DEPICT's call of PLINK. We used all defaults of DEPICT
# release 194. At this point we only used DEPICT to identify lead SNPs, not to perform
# biological annotation.
N_lead_SNPs_naive <- read.table(paste(
  data_dir, "DEPICT_run7/Neuroticism_model_3fact_GWASofN_naive_1E-5.clumped", sep=""),
  header=TRUE)

GWAS <-  userGWAS(LDSC_output, subset(neuroticism_sumstats, SNP %in% N_lead_SNPs_naive$SNP), 
                  "DWLS", model_3fact_GWASofN, modelchi=TRUE, printwarn=FALSE,
                  sub=c("N~SNP"), parallel=TRUE)
filename <- paste(data_dir, "Neuroticism_model_3fact_GWASofN_NleadSNPnaiveu_chiTRUE.txt", sep="")
write.table(GWAS, filename, sep="\t", row.names=FALSE, quote=FALSE)

GWAS <-  userGWAS(LDSC_output, subset(neuroticism_sumstats, SNP %in% N_lead_SNPs_naive$SNP), 
                  "DWLS", model_3fact_GWASofF, modelchi=TRUE, printwarn=FALSE,
                  sub=c("F1~SNP", "F2~SNP", "F3~SNP"), parallel=TRUE)
filename <- paste(data_dir, "Neuroticism_model_3fact_GWASofF_NleadSNPnaive_chiTRUE.txt", sep="")
write.table(GWAS, filename, sep="\t", row.names=FALSE, quote=FALSE)

GWAS <-  userGWAS(LDSC_output, subset(neuroticism_sumstats, SNP %in% N_lead_SNPs_naive$SNP), 
                  "DWLS", model_3fact_GWASindpath, modelchi=TRUE, printwarn=FALSE,
                  sub=c("mood~SNP", "misery~SNP", "irrit~SNP", "fedup~SNP", "lonely~SNP",
                        "nervous~SNP", "worry~SNP", "tense~SNP", "nerves~SNP", "guilt~SNP",
                        "hurt~SNP", "embarass~SNP"), parallel=TRUE)
filename <- paste(data_dir, "Neuroticism_model_3fact_GWASindpath_NleadSNPnaive_chiTRUE.txt", sep="")
write.table(GWAS, filename, sep="\t", row.names=FALSE, quote=FALSE)

# The above output file
# Neuroticism_model_3fact_GWASofN_NleadSNPnaive_chiTRUE.txt
# contains the lead SNPs from the GWAS of the Neuroticism general factor, as identified by
# DEPICT/PLINK clumping. Included are not only the GWAS results but also the model chi-square and
# degrees of freedom (as well as the AIC). The next output file contains the same SNPs but this
# time subjected to a GWAS (well, not literally genome wide) of the three group factors. The 
# numerical results are always given for the factors in the order Depressed Affect, Worry, and F3
# (recall that F3 has a loading on the general factor of 0.97). The third and final output file
# contains the same SNPs but this time subjected to GWAS specifying the independent-pathway model.

### END ###
####################################################################################################

setwd(data_dir)

N_lead_SNPs_naive_model1 <- read.delim(
  "Neuroticism_model_3fact_GWASofN_NleadSNPnaive_chiTRUE.txt"
)

N_lead_SNPs_naive_model2 <- read.delim(
  "Neuroticism_model_3fact_GWASofF_NleadSNPnaive_chiTRUE.txt"
)

N_lead_SNPs_naive_model3 <- read.delim(
  "Neuroticism_model_3fact_GWASindpath_NleadSNPnaive_chiTRUE.txt"
)

Q_chisq <- N_lead_SNPs_naive_model1$chisq - 
  N_lead_SNPs_naive_model2$chisq

# Some of the Q statistics are negative. Andrew Grotzinger has informed me (October 6, 2020)
# that this is a numerical problem that can happen when fits of the two models are very close.
# If this is the explanation, then a SNP's three path coefficients in model 2 should be 
# proportional to the SNP's one path coefficent in model 1 times the three factor loadings.

# The factor loadings of F on N are 0.796, 0.863, and 0.969.

SNP_pathcoef_mod1_F1 <- N_lead_SNPs_naive_model1$est*0.796
SNP_pathcoef_mod1_F2 <- N_lead_SNPs_naive_model1$est*0.863
SNP_pathcoef_mod1_F3 <- N_lead_SNPs_naive_model1$est*0.969

SNP_pathcoef_mod2_F1 <- N_lead_SNPs_naive_model2$est
SNP_pathcoef_mod2_F2 <- N_lead_SNPs_naive_model2$est.1
SNP_pathcoef_mod2_F3 <- N_lead_SNPs_naive_model2$est.2

cor.test(SNP_pathcoef_mod1_F1[which(Q_chisq<0)], SNP_pathcoef_mod2_F1[which(Q_chisq<0)])
cor.test(SNP_pathcoef_mod1_F2[which(Q_chisq<0)], SNP_pathcoef_mod2_F2[which(Q_chisq<0)])
cor.test(SNP_pathcoef_mod1_F3[which(Q_chisq<0)], SNP_pathcoef_mod2_F3[which(Q_chisq<0)])

cor.test(SNP_pathcoef_mod1_F1[which(Q_chisq>20)], SNP_pathcoef_mod2_F1[which(Q_chisq>20)])
cor.test(SNP_pathcoef_mod1_F2[which(Q_chisq>20)], SNP_pathcoef_mod2_F2[which(Q_chisq>20)])
cor.test(SNP_pathcoef_mod1_F3[which(Q_chisq>20)], SNP_pathcoef_mod2_F3[which(Q_chisq>20)])

# Correlations are greater than 0.99 when Q is negative, between 0.92 and 0.97 when Q is
# greater than 20.

df <- N_lead_SNPs_naive_model1$chisq_df - 
  N_lead_SNPs_naive_model2$chisq_df

Q_chisq_pval <- pchisq(Q_chisq, df, lower.tail=FALSE)

# We have calculated the Q_SNP statistic described by Grotzinger et al. (2019) for testing the
# significance of whether the SNP fits the simpler model. One problem with this approach is
# how to decide the cutoff. Should it really be P < 5E-8? In the future, as sample size
# increases even more, it may be that "too many" SNPs become significant.

# Originally we chose the model with the lowest AIC. A reviewer suggested Akaike weights,
# which seems reasonable.

get_AIC_weight <- function(AIC){
  delta_AIC <- AIC - min(AIC)
  den <- exp(-(1/2)*delta_AIC)
  output <- den/(sum(den)) 
  return(output) }

AIC_weight <- matrix(NA, nrow(N_lead_SNPs_naive_model1), ncol=3)
for(i in 1:nrow(AIC_weight)){
  AIC1 <- N_lead_SNPs_naive_model1[i,"AIC"]
  AIC2 <- N_lead_SNPs_naive_model2[i,"AIC"]
  AIC3 <- N_lead_SNPs_naive_model3[i,"AIC"]
  AIC_weight[i,] <- get_AIC_weight(c(AIC1, AIC2, AIC3)) }

# Say that a model wins if it gets more than 2/3.

# Write out the SNPs to files.

AIC_weight <- data.frame(AIC_weight)
colnames(AIC_weight) <- c("AIC1", "AIC2", "AIC3")

out <- data.frame(N_lead_SNPs_naive_model1, AIC_weight$AIC1)
colnames(out)[ncol(out)] <- "AIC1"
write.table(out[which(out$AIC1>(2/3)),],
            "N_lead_SNPs_AIC1.txt", quote=FALSE, row.names=FALSE, sep="\t")
            
out <- data.frame(N_lead_SNPs_naive_model2, AIC_weight$AIC2)
colnames(out)[ncol(out)] <- "AIC2"
write.table(out[which(out$AIC2>(2/3)),],
            "N_lead_SNPs_AIC2.txt", quote=FALSE, row.names=FALSE, sep="\t")            
            
out <- data.frame(N_lead_SNPs_naive_model3, AIC_weight$AIC3)
colnames(out)[ncol(out)] <- "AIC3"
write.table(out[which(out$AIC3>(2/3)),],
            "N_lead_SNPs_AIC3.txt", quote=FALSE, row.names=FALSE, sep="\t")
            
            
# The above output file
# N_lead_SNPs_AIC1.txt
# contains the lead SNPs from the GWAS of the Neuroticism general factor that have an Akaike
# weight exceeding 2/3, when compared with the group-factor (i.e., paths to the 3 group factors)
# and independent-pathway models. Those SNPs with Akaike weights exceeding 2/3 for one of
# the latter two models can be found in the AIC2 and AIC3 output files. Any SNP in the
# NleadSNPnaive_chiTRUE files that is not in one of these three AIC files does not have any
# Akaike weight exceeding 2/3.

###

# Now do GWAS of the group factors. Of the lead SNPs for a given factor, keep those that are not
# general-factor or independent-pathway SNPs from the general-factor GWAS and that have zero
# effects on the other group factors (as indicated by either chi-square-difference test or
# AIC weights).

####################################################################################################
### START ###

# Do not run the code in this block interactively. This will probably crash your GUI. Instead
# insert this block into a separate .R file and call it at the command line with Rscript.

require(GenomicSEM)

GWAS_dir <- # THE LOCATION OF THE NEUROTICISM SUMMARY STATS AT THE ITEM LEVEL
data_dir <- # THE LOCATION WHERE YOU WILL PLACE OUTPUT FILES
PLINK_call <- # THE LOCATION OF YOUR COPY OF PLINK 1.9

load(paste(data_dir, "Neuroticism_items_LDSC_output.Rdata", sep=""))
load(paste(data_dir, "Neuroticism_items_gSEM_sumstats.Rdata", sep=""))

model_3fact_GWASofF <- "F1 =~ mood + misery + irrit + fedup + lonely 
                        F2 =~ nervous + worry + tense + nerves
                        F3 =~ guilt + hurt + embarass
                        N =~ F1 + F2 + F3
                        F1 + F2 + F3 ~ SNP"

for(i in 1:22){
  print(i)
  GWAS <-  userGWAS(LDSC_output, subset(neuroticism_sumstats, CHR==i), "DWLS", model_3fact_GWASofF,
                    modelchi=FALSE, printwarn=FALSE,
                    sub=c("F1~SNP", "F2~SNP", "F3~SNP"), parallel=TRUE)
  filename <- paste(data_dir, "Neuroticism_model_3fact_GWASofF_chr", i, ".txt", sep="")
  write.table(GWAS, filename, sep="\t", row.names=FALSE, quote=FALSE)
  rm(GWAS)
  command <- paste("gzip ", filename, sep="")
  system(command) }

# WARNING
# The above loop might lead to exhaustion of memory, particularly after a larger chromosome.
# You might have to restart your machine if this happens. If you cannot execute this job on your
# local machine, please turn to whatever supercomputing resources are available to you.

setwd(data_dir)

# Insert the header into a new file that will contain the GWAS results from all chromosomes.
command <- paste("gzcat Neuroticism_model_3fact_GWASofF_chr22.txt.gz | ",
                 "head -n 1 > ",
                 "Neuroticism_model_3fact_GWASofN_allchr.txt",
                 sep="")
system(command)

for(i in 1:22){
  print(i)
  command <- paste("gzcat Neuroticism_model_3fact_GWASofF_chr", i, ".txt.gz | ",
                   "tail -n +2 >> ",
                   "Neuroticism_model_3fact_GWASofF_allchr.txt",
                   sep="")
  system(command) }

system("gzip Neuroticism_model_3fact_GWASofF_allchr.txt")

### END ###
####################################################################################################

# This file .clumped contains lead SNPs in the GWAS of the Depressed Affect group factor
# (model_3fact_GWASofF1), as identified by DEPICT's call of PLINK. We used all defaults of DEPICT
# release 194. At this point we only used DEPICT to identify lead SNPs, not to perform
# biological annotation.
F1_lead_SNPs_naive <- read.table(paste(
  data_dir, "Neuroticism_model_3fact_GWASofF1_naive_1E-5.clumped", sep=""),
  header=TRUE)

# Get rid of highly weighted lead SNPs for Neuroticism or independent pathways from earlier.
N_lead_SNPs_AIC1 <- read.delim("N_lead_SNPs_AIC1.txt")
N_lead_SNPs_AIC3 <- read.delim("N_lead_SNPs_AIC3.txt")
F1_lead_SNPs <- subset(F1_lead_SNPs_naive,
                       !(SNP %in% N_lead_SNPs_AIC1$SNP | SNP %in% N_lead_SNPs_AIC3$SNP))
write.table(F1_lead_SNPs, "F1_lead_SNPs.txt", sep="\t", row.names=FALSE, quote=FALSE)

####################################################################################################
### START ###

require(GenomicSEM)

GWAS_dir <- # THE LOCATION OF THE NEUROTICISM SUMMARY STATS AT THE ITEM LEVEL
data_dir <- # THE LOCATION WHERE YOU WILL PLACE OUTPUT FILES
PLINK_call <- # THE LOCATION OF YOUR COPY OF PLINK 1.9

load(paste(data_dir, "Neuroticism_items_LDSC_output.Rdata", sep=""))
load(paste(data_dir, "Neuroticism_items_gSEM_sumstats.Rdata", sep=""))

model_3fact_GWASofN <-"F1 =~ mood + misery + irrit + fedup + lonely 
                       F2 =~ nervous + worry + tense + nerves
                       F3 =~ guilt + hurt + embarass
                        N =~ F1 + F2 + F3
                        N ~ SNP"

model_3fact_GWASofF <- "F1 =~ mood + misery + irrit + fedup + lonely 
                        F2 =~ nervous + worry + tense + nerves
                        F3 =~ guilt + hurt + embarass
                        N =~ F1 + F2 + F3
                        F1 + F2 + F3 ~ SNP"

model_3fact_GWASofF1 <- "F1 =~ mood + misery + irrit + fedup + lonely 
                         F2 =~ nervous + worry + tense + nerves
                         F3 =~ guilt + hurt + embarass
                         N =~ F1 + F2 + F3
                         F1 ~ SNP"

model_3fact_GWASindpath <- "F1 =~ mood + misery + irrit + fedup + lonely 
                        F2 =~ nervous + worry + tense + nerves
                        F3 =~ guilt + hurt + embarass
                        N =~ F1 + F2 + F3
mood + misery + irrit + fedup + lonely + nervous + worry + tense + nerves + guilt + hurt + embarass ~ SNP"


F1_lead_SNPs <- read.delim(paste(
  data_dir, "F1_lead_SNPs.txt", sep=""))

# GWAS <-  userGWAS(LDSC_output, subset(neuroticism_sumstats, SNP %in% F1_lead_SNPs$SNP), 
                  # "DWLS", model_3fact_GWASofN, modelchi=TRUE, printwarn=FALSE,
                  # sub=c("N~SNP"), parallel=TRUE)
# filename <- paste(data_dir, "Neuroticism_model_3fact_GWASofN_F1leadSNP_chiTRUE.txt", sep="")
# write.table(GWAS, filename, sep="\t", row.names=FALSE, quote=FALSE)

GWAS <-  userGWAS(LDSC_output, subset(neuroticism_sumstats, SNP %in% F1_lead_SNPs$SNP), 
                  "DWLS", model_3fact_GWASofF, modelchi=TRUE, printwarn=FALSE,
                  sub=c("F1~SNP", "F2~SNP", "F3~SNP"), parallel=TRUE)
filename <- paste(data_dir, "Neuroticism_model_3fact_GWASofF_F1leadSNP_chiTRUE.txt", sep="")
write.table(GWAS, filename, sep="\t", row.names=FALSE, quote=FALSE)

GWAS <-  userGWAS(LDSC_output, subset(neuroticism_sumstats, SNP %in% F1_lead_SNPs$SNP), 
                  "DWLS", model_3fact_GWASofF1, modelchi=TRUE, printwarn=FALSE,
                  sub=c("F1~SNP"), parallel=TRUE)
filename <- paste(data_dir, "Neuroticism_model_3fact_GWASofF1_F1leadSNP_chiTRUE.txt", sep="")
write.table(GWAS, filename, sep="\t", row.names=FALSE, quote=FALSE)

GWAS <-  userGWAS(LDSC_output, subset(neuroticism_sumstats, SNP %in% F1_lead_SNPs$SNP), 
                  "DWLS", model_3fact_GWASindpath, modelchi=TRUE, printwarn=FALSE,
                  sub=c("mood~SNP", "misery~SNP", "irrit~SNP", "fedup~SNP", "lonely~SNP",
                        "nervous~SNP", "worry~SNP", "tense~SNP", "nerves~SNP", "guilt~SNP",
                        "hurt~SNP", "embarass~SNP"), parallel=TRUE)
filename <- paste(data_dir, "Neuroticism_model_3fact_GWASindpath_F1leadSNP_chiTRUE.txt", sep="")
write.table(GWAS, filename, sep="\t", row.names=FALSE, quote=FALSE)

# The above output file
# Neuroticism_model_3fact_GWASofF_F1leadSNPnaive_chiTRUE.txt
# contains the lead SNPs from the GWAS of Depressed Affect, as identified by
# DEPICT/PLINK clumping. Included are not only the GWAS results but also the model chi-square and
# degrees of freedom (as well as the AIC). The next output file contains the same SNPs but this
# time put through a Genomic SEM run setting the paths to the other two
# group factors to zero. The third and final output file contains the same SNPs but this time 
# put through a Genomic SEM run specifying the independent-pathway model.

### END ###
####################################################################################################

setwd(data_dir)

F1_lead_SNPs_model1 <- read.delim(
  "Neuroticism_model_3fact_GWASofF1_F1leadSNP_chiTRUE.txt"
)

F1_lead_SNPs_model2 <- read.delim(
  "Neuroticism_model_3fact_GWASofF_F1leadSNP_chiTRUE.txt"
)

F1_lead_SNPs_model3 <- read.delim(
  "Neuroticism_model_3fact_GWASindpath_F1leadSNP_chiTRUE.txt"
)

Q_chisq <- F1_lead_SNPs_model1$chisq - 
  F1_lead_SNPs_model2$chisq
# This time they are all positive. It seems not that many SNPs only affect F1.

df <- F1_lead_SNPs_model1$chisq_df - 
  F1_lead_SNPs_model2$chisq_df
  
Q_chisq_pval <- pchisq(Q_chisq, df, lower.tail=FALSE)

F1_lead_SNPs_model2[which(Q_chisq_pval>.05),c("Z_Estimate", "Z_Estimate.1", "Z_Estimate.2")]
# Yes, Z stat of F1 is larger in magnitude than those of F2 and F3.

AIC_weight <- matrix(NA, nrow(F1_lead_SNPs_model1), ncol=3)
for(i in 1:nrow(AIC_weight)){
  AIC1 <- F1_lead_SNPs_model1[i,"AIC"]
  AIC2 <- F1_lead_SNPs_model2[i,"AIC"]
  AIC3 <- F1_lead_SNPs_model3[i,"AIC"]
  AIC_weight[i,] <- get_AIC_weight(c(AIC1, AIC2, AIC3)) }

# There are hardly any SNPs where an effect only on F1 has an Akaike weight > (2/3).

# Write out the SNPs to files.
            
AIC_weight <- data.frame(AIC_weight)
colnames(AIC_weight) <- c("AIC1", "AIC2", "AIC3")

out <- data.frame(F1_lead_SNPs_model1, AIC_weight$AIC1)
colnames(out)[ncol(out)] <- "AIC1"
write.table(out[which(out$AIC1>(2/3)),],
            "F1_lead_SNPs_AIC1.txt", quote=FALSE, row.names=FALSE, sep="\t")
            
out <- data.frame(F1_lead_SNPs_model2, AIC_weight$AIC2)
colnames(out)[ncol(out)] <- "AIC2"
write.table(out[which(out$AIC2>(2/3)),],
            "F1_lead_SNPs_AIC2.txt", quote=FALSE, row.names=FALSE, sep="\t")            
            
out <- data.frame(F1_lead_SNPs_model3, AIC_weight$AIC3)
colnames(out)[ncol(out)] <- "AIC3"
write.table(out[which(out$AIC3>(2/3)),],
            "F1_lead_SNPs_AIC3.txt", quote=FALSE, row.names=FALSE, sep="\t")            

# The above output file
# F1_lead_SNPs_AIC1.txt
# contains the lead SNPs from the GWAS of Depressed Affect that have an Akaike
# weight exceeding 2/3 in favor of a path to Depressed Affect only, when compared with the 
# group-factor (i.e., paths to the 3 group factors) and independent-pathway models. Those SNPs
# with Akaike weights exceeding 2/3 for one of
# the latter two models can be found in the AIC2 and AIC3 output files. Any SNP in the
# F1leadSNPnaive_chiTRUE files that is not in one of these three AIC files does not have any
# Akaike weight exceeding 2/3.

### Now for F2.
# All output files are closely analagous to those for F1 above.

F2_lead_SNPs_naive <- read.table(paste(
  data_dir, "Neuroticism_model_3fact_GWASofF2_naive_1E-5.clumped", sep=""),
  header=TRUE)

# Get rid of highly weighted lead SNPs for Neuroticism, F1, or independent pathways from earlier.
N_lead_SNPs_AIC1 <- read.delim("N_lead_SNPs_AIC1.txt")
N_lead_SNPs_AIC3 <- read.delim("N_lead_SNPs_AIC3.txt")
F1_lead_SNPs_AIC1 <- read.delim("F1_lead_SNPs_AIC1.txt")
F1_lead_SNPs_AIC2 <- read.delim("F1_lead_SNPs_AIC2.txt")
F1_lead_SNPs_AIC3 <- read.delim("F1_lead_SNPs_AIC3.txt")
F2_lead_SNPs <- subset(F2_lead_SNPs_naive,
                       !(SNP %in% N_lead_SNPs_AIC1$SNP | SNP %in% N_lead_SNPs_AIC3$SNP |
                         SNP %in% F1_lead_SNPs_AIC1$SNP | SNP %in% F1_lead_SNPs_AIC2$SNP |
                         SNP %in% F1_lead_SNPs_AIC3$SNP))
write.table(F2_lead_SNPs, "F2_lead_SNPs.txt", sep="\t", row.names=FALSE, quote=FALSE)

####################################################################################################
### START ###

require(GenomicSEM)

GWAS_dir <- # THE LOCATION OF THE NEUROTICISM SUMMARY STATS AT THE ITEM LEVEL
data_dir <- # THE LOCATION WHERE YOU WILL PLACE OUTPUT FILES
PLINK_call <- # THE LOCATION OF YOUR COPY OF PLINK 1.9

load(paste(data_dir, "Neuroticism_items_LDSC_output.Rdata", sep=""))
load(paste(data_dir, "Neuroticism_items_gSEM_sumstats.Rdata", sep=""))

model_3fact_GWASofN <-"F1 =~ mood + misery + irrit + fedup + lonely 
                       F2 =~ nervous + worry + tense + nerves
                       F3 =~ guilt + hurt + embarass
                        N =~ F1 + F2 + F3
                        N ~ SNP"

model_3fact_GWASofF <- "F1 =~ mood + misery + irrit + fedup + lonely 
                        F2 =~ nervous + worry + tense + nerves
                        F3 =~ guilt + hurt + embarass
                        N =~ F1 + F2 + F3
                        F1 + F2 + F3 ~ SNP"

model_3fact_GWASofF2 <- "F1 =~ mood + misery + irrit + fedup + lonely 
                         F2 =~ nervous + worry + tense + nerves
                         F3 =~ guilt + hurt + embarass
                         N =~ F1 + F2 + F3
                         F2 ~ SNP"

model_3fact_GWASindpath <- "F1 =~ mood + misery + irrit + fedup + lonely 
                        F2 =~ nervous + worry + tense + nerves
                        F3 =~ guilt + hurt + embarass
                        N =~ F1 + F2 + F3
mood + misery + irrit + fedup + lonely + nervous + worry + tense + nerves + guilt + hurt + embarass ~ SNP"


F2_lead_SNPs <- read.delim(paste(
  data_dir, "F2_lead_SNPs.txt", sep=""))

GWAS <-  userGWAS(LDSC_output, subset(neuroticism_sumstats, SNP %in% F2_lead_SNPs$SNP), 
                  "DWLS", model_3fact_GWASofN, modelchi=TRUE, printwarn=FALSE,
                  sub=c("N~SNP"), parallel=TRUE)
filename <- paste(data_dir, "Neuroticism_model_3fact_GWASofN_F2leadSNP_chiTRUE.txt", sep="")
write.table(GWAS, filename, sep="\t", row.names=FALSE, quote=FALSE)

GWAS <-  userGWAS(LDSC_output, subset(neuroticism_sumstats, SNP %in% F2_lead_SNPs$SNP), 
                  "DWLS", model_3fact_GWASofF, modelchi=TRUE, printwarn=FALSE,
                  sub=c("F1~SNP", "F2~SNP", "F3~SNP"), parallel=TRUE)
filename <- paste(data_dir, "Neuroticism_model_3fact_GWASofF_F2leadSNP_chiTRUE.txt", sep="")
write.table(GWAS, filename, sep="\t", row.names=FALSE, quote=FALSE)

GWAS <-  userGWAS(LDSC_output, subset(neuroticism_sumstats, SNP %in% F2_lead_SNPs$SNP), 
                  "DWLS", model_3fact_GWASofF2, modelchi=TRUE, printwarn=FALSE,
                  sub=c("F2~SNP"), parallel=TRUE)
filename <- paste(data_dir, "Neuroticism_model_3fact_GWASofF2_F2leadSNP_chiTRUE.txt", sep="")
write.table(GWAS, filename, sep="\t", row.names=FALSE, quote=FALSE)

GWAS <-  userGWAS(LDSC_output, subset(neuroticism_sumstats, SNP %in% F2_lead_SNPs$SNP), 
                  "DWLS", model_3fact_GWASindpath, modelchi=TRUE, printwarn=FALSE,
                  sub=c("mood~SNP", "misery~SNP", "irrit~SNP", "fedup~SNP", "lonely~SNP",
                        "nervous~SNP", "worry~SNP", "tense~SNP", "nerves~SNP", "guilt~SNP",
                        "hurt~SNP", "embarass~SNP"), parallel=TRUE)
filename <- paste(data_dir, "Neuroticism_model_3fact_GWASindpath_F2leadSNP_chiTRUE.txt", sep="")
write.table(GWAS, filename, sep="\t", row.names=FALSE, quote=FALSE)

### END ###
####################################################################################################

setwd(data_dir)

F2_lead_SNPs_model1 <- read.delim(
  "Neuroticism_model_3fact_GWASofF2_F2leadSNP_chiTRUE.txt"
)

F2_lead_SNPs_model2 <- read.delim(
  "Neuroticism_model_3fact_GWASofF_F2leadSNP_chiTRUE.txt"
)

F2_lead_SNPs_model3 <- read.delim(
  "Neuroticism_model_3fact_GWASindpath_F2leadSNP_chiTRUE.txt"
)

Q_chisq <- F2_lead_SNPs_model1$chisq - 
  F2_lead_SNPs_model2$chisq
# All positive. It seem not that many SNPs only affect F2.

df <- F2_lead_SNPs_model1$chisq_df - 
  F2_lead_SNPs_model2$chisq_df

Q_chisq_pval <- pchisq(Q_chisq, df, lower.tail=FALSE)

F2_lead_SNPs_model2[which(Q_chisq_pval>.05),c("Z_Estimate", "Z_Estimate.1", "Z_Estimate.2")]
# Yes, Z stat of F2 is larger in magnitude than those of F1 and F3.

AIC_weight <- matrix(NA, nrow(F2_lead_SNPs_model1), ncol=3)
for(i in 1:nrow(AIC_weight)){
  AIC1 <- F2_lead_SNPs_model1[i,"AIC"]
  AIC2 <- F2_lead_SNPs_model2[i,"AIC"]
  AIC3 <- F2_lead_SNPs_model3[i,"AIC"]
  AIC_weight[i,] <- get_AIC_weight(c(AIC1, AIC2, AIC3)) }

# There are hardly any SNPs where an effect only on F2 has an Akaike weight > (2/3).

# Write out the SNPs to files.

AIC_weight <- data.frame(AIC_weight)
colnames(AIC_weight) <- c("AIC1", "AIC2", "AIC3")

out <- data.frame(F2_lead_SNPs_model1, AIC_weight$AIC1)
colnames(out)[ncol(out)] <- "AIC1"
write.table(out[which(out$AIC1>(2/3)),],
            "F2_lead_SNPs_AIC1.txt", quote=FALSE, row.names=FALSE, sep="\t")
            
out <- data.frame(F2_lead_SNPs_model2, AIC_weight$AIC2)
colnames(out)[ncol(out)] <- "AIC2"
write.table(out[which(out$AIC2>(2/3)),],
            "F2_lead_SNPs_AIC2.txt", quote=FALSE, row.names=FALSE, sep="\t")            
            
out <- data.frame(F2_lead_SNPs_model3, AIC_weight$AIC3)
colnames(out)[ncol(out)] <- "AIC3"
write.table(out[which(out$AIC3>(2/3)),],
            "F2_lead_SNPs_AIC3.txt", quote=FALSE, row.names=FALSE, sep="\t")

# To do biological annotation of independent-pathway SNPs, just those SNPs from the above
# analyses that have big Akaike weights for the independent-pathway model. Just need their
# SNP, CHR, BP, and p-value (which I can make up).

N_indpath <- read.delim("N_lead_SNPs_AIC3.txt")[,c("SNP", "CHR", "BP")]
F1_indpath <- read.delim("F1_lead_SNPs_AIC3.txt")[,c("SNP", "CHR", "BP")]
F2_indpath <- read.delim("F2_lead_SNPs_AIC3.txt")[,c("SNP", "CHR", "BP")]
indpath <- rbind(N_indpath, F1_indpath, F2_indpath)
indpath <- data.frame(indpath, 1e-6)
colnames(indpath)[4] <- "P"
write.table(indpath, "indpath_lead_SNPs.txt", quote=FALSE, row.names=FALSE, sep="\t")

# How much do the general-factor and independent-pathway loci overlap?

N_genes <- read.delim("Neuroticism_model_3fact_GWASofN_AIC1_1E-5_geneprioritization.txt",
                      header=TRUE)
indpath_genes <- read.delim("Neuroticism_model_3fact_indpath_geneprioritization.txt") 

intersect(N_genes$Gene.symbol, indpath_genes$Gene.symbol)    

# It is tedious to make the supplementary table of independent-pathway SNPs by hand. Do it here.

N_indpath <- read.delim("N_lead_SNPs_AIC3.txt")
F1_indpath <- read.delim("F1_lead_SNPs_AIC3.txt")
F2_indpath <- read.delim("F2_lead_SNPs_AIC3.txt")
indpath <- rbind(N_indpath, F1_indpath, F2_indpath)

# To calculate Q_SNP, I need to get the model2 chiTRUE GWAS of these SNPs.

N_lead_SNPs_model2 <- read.delim(
  "Neuroticism_model_3fact_GWASofF_NleadSNPnaive_chiTRUE.txt"
)

F1_lead_SNPs_model2 <- read.delim(
  "Neuroticism_model_3fact_GWASofF_F1leadSNP_chiTRUE.txt"
)

F2_lead_SNPs_model2 <- read.delim(
  "Neuroticism_model_3fact_GWASofF_F2leadSNP_chiTRUE.txt"
)
               
lead_SNPs_model2 <- rbind(N_lead_SNPs_model2, F1_lead_SNPs_model2, F2_lead_SNPs_model2)               
indpath <- merge(indpath, lead_SNPs_model2[,c("SNP", "chisq", "chisq_df")], by="SNP", all.x=TRUE)
indpath <- indpath[!duplicated(indpath),]

Q_chisq <- indpath$chisq.y - indpath$chisq.x
df <- indpath$chisq_df.y - indpath$chisq_df.x
Q_chisq_pval <- pchisq(Q_chisq, df, lower.tail=FALSE)

indpath <- data.frame(indpath, Q_chisq, Q_chisq_pval)

out <- indpath[,c(
"SNP",
"CHR",
"BP",
"MAF",
"A1",
"A2",
"est",
"SE",
"Pval_Estimate",
"est.1",
"SE.1",
"Pval_Estimate.1",
"est.2",
"SE.2",
"Pval_Estimate.2",
"est.3",
"SE.3",
"Pval_Estimate.3",
"est.4",
"SE.4",
"Pval_Estimate.4",
"est.5",
"SE.5",
"Pval_Estimate.5",
"est.6",
"SE.6",
"Pval_Estimate.6",
"est.7",
"SE.7",
"Pval_Estimate.7",
"est.8",
"SE.8",
"Pval_Estimate.8",
"est.9",
"SE.9",
"Pval_Estimate.9",
"est.10",
"SE.10",
"Pval_Estimate.10",
"est.11",
"SE.11",
"Pval_Estimate.11",
"Q_chisq",
"Q_chisq_pval",
"AIC3"
)]

colnames(out) <- c(
"SNP",
"CHR",
"BP",
"MAF",
"A1",
"A2",
"Mood effect",
"Mood SE",
"Mood GWAS p value",
"Misery effect",
"Misery SE",
"Misery GWAS p value",
"Irrit effect",
"Irrit SE",
"Irrit GWAS p value",
"Hurt effect",
"Hurt SE",
"Hurt GWAS pvalue",
"Fed-up effect",
"Fed-up SE",
"Fed-up GWAS p value",
"Nervous effect",
"Nervous SE",
"Nervous GWAS p value",
"Worry effect",
"Worry SE",
"Worry GWAS p value",
"Tense effect",
"Tense SE",
"Tense GWAS p value",
"Embarass effect",
"Embarass SE",
"Embarass GWAS p value",
"Nerves effect",
"Nerves SE",
"Nerves GWAS p value",
"Lonely effect",
"Lonely SE",
"Lonely GWAS p value",
"Guilt effect",
"Guilt SE",
"Guilt GWAS p value",
"Q_SNP",
"Q_SNP p value",
"Akaike weight"
)

write.table(out, "all_indpath_SNPs.txt", sep="\t", row.names=FALSE, quote=FALSE)
# This is essentially the same as Supplementary Table S10.  

# How much sign concordance?
indpath <- read.delim("all_indpath_SNPs.txt")   

concordance <- rep(NA, nrow(indpath))
for(i in 1:nrow(indpath)){
	print(i)
	pos_effect <- indpath[i,c("Mood.effect", "Misery.effect", "Irrit.effect",
	                          "Fed.up.effect", "Nervous.effect", "Worry.effect",
	                          "Tense.effect", "Embarass.effect", "Nerves.effect",
	                          "Lonely.effect", "Guilt.effect", "Guilt.effect")] > 0
	concord <- sum(pos_effect)
	if(concord < 6) concord <- 12 - concord
	concordance[i] <- concord }      

####################################################################################################

# To calculate the genetic correlations with the residuals of depressed affect and worry, it will
# perhaps to be simplest to switch to a bifactor model so that the group factors already have no
# loadings on the general factor.

model_bifact <- "F1 =~ mood + misery + irrit + fedup + lonely 
                 F2 =~ nervous + worry + tense + nerves
                 N =~ mood + misery + irrit + fedup + lonely +
                      nervous + worry + tense + nerves + 
                      guilt + hurt + embarass
                 N ~~ 0*F1
                 N ~~ 0*F2
                 F1 ~~ 0*F2
                 nervous ~~ 0*nervous"
                 
model_bifact_out <- usermodel(LDSC_output, "DWLS", model_bifact, CFIcalc=TRUE, std.lv=FALSE,
                              imp_cov=TRUE)
# Fit is slightly better than hierarchical. Some difficulties in fitting the model. Initially
# I included the factor F3, but this led to a convergence time of ~45 seconds (unacceptable for
# a GWAS over ~7M SNPs) and Heywood cases. Dropping F3 led to a convergence time similar to that 
# of the hierarchical model. I fixed the residual variance of nervous to zero to prevent it from
# becoming negative.

####################################################################################################
### START ###

# Do not run the code in this block interactively. This will probably crash your GUI. Instead
# insert this block into a separate .R file and call it at the command line with Rscript.

require(GenomicSEM)

GWAS_dir <- # THE LOCATION OF THE NEUROTICISM SUMMARY STATS AT THE ITEM LEVEL
data_dir <- # THE LOCATION WHERE YOU WILL PLACE OUTPUT FILES
PLINK_call <- # THE LOCATION OF YOUR COPY OF PLINK 1.9

load(paste(data_dir, "Neuroticism_items_LDSC_output.Rdata", sep=""))
load(paste(data_dir, "Neuroticism_items_gSEM_sumstats.Rdata", sep=""))

model_bifact_GWASofN <- "F1 =~ mood + misery + irrit + fedup + lonely 
                         F2 =~ nervous + worry + tense + nerves
                         N =~ mood + misery + irrit + fedup + lonely +
                              nervous + worry + tense + nerves + 
                              guilt + hurt + embarass
                         N ~~ 0*F1
                         N ~~ 0*F2
                         F1 ~~ 0*F2
                         nervous ~~ 0*nervous
                         N + F1 + F2 ~ SNP"

for(i in 1:22){
  print(i)
  GWAS <-  userGWAS(LDSC_output, subset(neuroticism_sumstats, CHR==i), "DWLS", model_bifact_GWASofN,
                    printwarn=TRUE, sub=c("N~SNP", "F1~SNP", "F2~SNP"), parallel=TRUE,
                    toler=1e-30)
                    # In one run I got a warning about the tolerance for matrix inversion. 
                    # Hence the change.                    
  filename <- paste(data_dir, "Neuroticism_model_bifact_GWASofN_chr", i, ".txt", sep="")
  write.table(GWAS, filename, sep="\t", row.names=FALSE, quote=FALSE)
  rm(GWAS)
  command <- paste("gzip ", filename, sep="")
  system(command) }

# WARNING
# The above loop might lead to exhaustion of memory, particularly after a larger chromosome.
# You might have to restart your machine if this happens. If you cannot execute this job on your
# local machine, please turn to whatever supercomputing resources are available to you.

setwd(data_dir)

# Insert the header into a new file that will contain the GWAS results from all chromosomes.
command <- paste("gzcat Neuroticism_model_bifact_GWASofN_chr22.txt.gz | ",
                 "head -n 1 > ",
                 "Neuroticism_model_bifact_GWASofN_allchr.txt",
                 sep="")
system(command)

for(i in 1:22){
  print(i)
  command <- paste("gzcat Neuroticism_model_bifact_GWASofN_chr", i, ".txt.gz | ",
                   "tail -n +2 >> ",
                   "Neuroticism_model_bifact_GWASofN_allchr.txt",
                   sep="")
  system(command) }

system("gzip Neuroticism_model_bifact_GWASofN_allchr.txt")

### END ###                                              
####################################################################################################

# The summary statistic generated above contain two additional groups of columns with the suffixes
# .1 and .2. These can be used to calculate genetic correlations with other traits.
```
