## Supplementary material for "Genetic and neural bases of the neuroticism general factor": README.txt

The file genomicSEM_revision_forpub.R contains more information about the files in this directory.

###

Neuroticism_model_3fact_GWASofG_NleadSNPnaive_chiTRUE.txt
Neuroticism_model_3fact_GWASofF_NleadSNPnaive_chiTRUE.txt
Neuroticism_model_3fact_GWASindpath_NleadSNPnaive_chiTRUE.txt

These three files contain Genomic SEM output for the same SNPs -- the lead SNPs in the GWAS of the Neuroticism general factor, identified by the DEPICT/PLINK clumping procedure. The tested path model encompassing the SNP differs across files. In the first file, the SNP sends a path to the general factor, as in the GWAS used to identify these SNPs. In the second file, the SNP sends paths to the three group factors--in order, Depressed Affect, Worry, and the factor that we have left unnamed because of its near-unity loading on the hierarchical second-order factor of Neuroticism. In the third file, the SNP sends paths to all 12 items ("independent pathways"). The files contain model chi-square statistics, degrees of freedom, and AICs, allowing the user to calculate the Q_SNP statistic of Grotzinger et al. (2019) and Akaike weights.

###

N_lead_SNPs_AIC1.txt
N_lead_SNPs_AIC2.txt
N_lead_SNPs_AIC3.txt

The first file in this set contains the lead SNPs from the GWAS of Neuroticism that attain an Akaike weight for the general-factor path model (SNP --> general factor) greater than 2/3. The second file contains those lead SNPs from the same GWAS that attain an Akaike weight for the group-factor model (SNP sends paths to the 3 group factors) greater than 2/3. The third file contains those lead SNPs from the same GWAS that attain an Akaike weight for the independent-pathway model greater than 2/3. SNPs failing to attain an Akaike weight greater than 2/3 for any model can still be found in the "NleadSNPnaive_chiTRUE" files listed above.

###

Neuroticism_model_3fact_GWASofF1_F1leadSNP_chiTRUE.txt
Neuroticism_model_3fact_GWASofF_F1leadSNP_chiTRUE.txt
Neuroticism_model_3fact_GWASindpath_F1leadSNP_chiTRUE.txt

These three files contain Genomic SEM output for the same SNPs -- the lead SNPs in the GWAS of Depressed Affect, identified by applying the DEPICT/PLINK clumping procedure to the Depressed Affect columns in the output of the GWAS of all three group factors (i.e., a path model sending arrows to all three group factors). The tested path model encompassing the SNP differs across files. In the first file, the SNP sends a path to Depressed Affect only. In the second file, the SNP sends paths to the three group factors, as in the GWAS used for discovery of these SNPs initially. The files contain model chi-square statistics, degrees of freedom, and AICs, allowing the user to calculate the Q_SNP statistic of Grotzinger et al. (2019) and Akaike weights.

###

F1_lead_SNPs_AIC1.txt
F1_lead_SNPs_AIC2.txt
F1_lead_SNPs_AIC3.txt

The first file in this set contains the lead SNPs from the GWAS of the three group factors that attain an Akaike weight for affecting Depressed Affect only greater than 2/3. The second file contains those lead SNPs from the same GWAS that attain an Akaike weight for the group-factor model (SNP sends paths to the 3 group factors) greater than 2/3. The third file contains those lead SNPs from the same GWAS that attain an Akaike weight for the independent-pathway model greater than 2/3. SNPs failing to attain an Akaike weight greater than 2/3 for any model can still be found in the "F1leadSNPnaive_chiTRUE" files listed above.

###

Neuroticism_model_3fact_GWASofF2_F2leadSNP_chiTRUE.txt
Neuroticism_model_3fact_GWASofF_F2leadSNP_chiTRUE.txt
Neuroticism_model_3fact_GWASindpath_F2leadSNP_chiTRUE.txt

F2_lead_SNPs_AIC1.txt
F2_lead_SNPs_AIC2.txt
F2_lead_SNPs_AIC3.txt


These files for the Worry group factor are analogous to their Depressed Affect counterparts above.
