## Supplementary material for "Genetic and neural bases of the neuroticism general factor": S-LDSC.R

```
####################################################################################################
library(RColorBrewer)

ldsc_path <- # THE DIRECTORY WHERE YOU KEEP LD SCORE REGRESSION (LDSC)
data_dir <- # THE DIRECTORY WHERE YOU WANT TO PLACE THE OUTPUT

setwd(data_dir)

# Munge the summary statistics.

# munge_sumstats.py is not working for some reason. Manually convert.

d <- read.delim("Neuroticism_model_3fact_GWASofN_allchr.txt.gz")
# If the summary statistics of the factor GWAS are not in data_dir, specify the path.

# columns required by LDSC are SNP, N, Z, A1, A2

N <- rep(375000, nrow(d))
# The precise sample size does not matter for Genomic SEM factors. 375,000 is close to the actual
# sample sizes of the item-level GWAS.

out <- data.frame(d$SNP, N, d$Z_Estimate, d$A1, d$A2)
colnames(out) <- c("SNP", "N", "Z", "A1", "A2")
write.table(out, "GWASofN_hierarchical.sumstats", quote=FALSE, row.names=FALSE, sep="\t")

# As a test, just do baseline.

command <- paste(ldsc_path, "ldsc.py",
                 " --h2 ", data_dir, "GWASofN_hierarchical.sumstats.gz",
                 " --ref-ld-chr ", ldsc_path, "files/LDSCORE_1000G_Phase3_baselineLD_v2.2_ldscores/",
                 "baselineLD.",
                 " --w-ld-chr ", ldsc_path, "files/weights_hm3_no_hla/weights.",
                 " --overlap-annot",
                 " --frqfile-chr ", ldsc_path, "files/1000G_Phase3_frq/1000G.EUR.QC.",
                 " --print-coefficients",
                 " --out ", data_dir, "baseline",
                 sep="") 
# My LDSC directory has a subdirectory called "files" with its own subdirectories at a deeper 
# level. Your setup may differ.                 

system(command)
                 
baseline <- read.delim("baseline.results")                 
# This is essentially Supplementary Table S15.

### heritability enrichment of DEPICT tissues

for(i in 1:152){
	command <- paste(ldsc_path, "ldsc.py",
                 " --h2 ", data_dir, "GWASofN_hierarchical.sumstats.gz",
                 " --ref-ld-chr ", ldsc_path, "files/1000G_Phase3_Franke/Franke.", 
                 i, ".,",
                 # I downloaded the Franke files just for this analysis.
                 ldsc_path, "files/1000G_Phase3_Franke/Franke.control.,",  # any-gene control              
                 ldsc_path, "files/LDSCORE_1000G_Phase3_baselineLD_v2.2_ldscores/baselineLD.",
                 " --w-ld-chr ", ldsc_path, "files/weights_hm3_no_hla/weights.",
                 " --overlap-annot",
                 " --frqfile-chr ", ldsc_path, "files/1000G_Phase3_frq/1000G.EUR.QC.",
                 " --print-coefficients",
                 " --out ", data_dir, "DEPICT_tissue_", i,
                 sep="") 
                 
    system(command)  }

d <- read.delim("DEPICT_tissue_1.results")
d <- d[1,]
for(i in 2:152){
	print(i)
	new <- read.delim(paste("DEPICT_tissue_", i, ".results", sep=""))
	d <- rbind(d, new[1,]) }
	
tissue_ID <- read.table(paste(ldsc_path, 
             "files/1000G_Phase3_Franke/Franke.cell_types.txt", sep="")	
d[,1] <- tissue_ID[,1]

# Finucane et al. (2018) filtered the DEPICT tissues to eliminate redundant or highly correlated
# tissues/cell types. Their criteria for which to retain were similar to that of Lee et al. 
# (2018). Create a vector of retained identifiers to subset the *_tissueenrichment.txt ouput
# of DEPICT.

keep <- c(
"Adrenal Cortex",
"Adrenal Glands",
"Aortic Valve",
"Arteries",
"B Lymphocytes",
"Basal Ganglia",
"Bone Marrow Cells",
"Blood Vessels",
"Blood",
"Blood Platelets",
"Blood Cells",
"Urinary Bladder",
"Brain Stem",
"Brain",
"Cerebellum",
"Cerebral Cortex",
"Cartilage",
"Cecum",
"Chorion",
"Cervical Vertebrae",
"Cervix Uteri",
"Colon",
"Cicatrix",
"Corpus Striatum",
"Connective Tissue Cells",
"Dentition",
"Dendritic Cells",
"Diencephalon",
"Epithelium",
"Epithelial Cells",
"Epidermis",
"Esophagus",
"Endometrium",
"Endocrine Glands",
"Frontal Lobe",
"Eye",
"Fallopian Tubes",
"Fetal Blood",
"Fibroblasts",
"Gonads",
"Granulosa Cells",
"Germ Cells",
"Genitalia",
"Genitalia  Female",
"Genitalia  Male",
"Hippocampus",
"Heart",
"Heart Atria",
"Heart Ventricles",
"Hematopoietic Stem Cells",
"Ileum",
"Immune System",
"Hypothalamus",
"Hypothalamo Hypophyseal System",
"Keloid",
"Kidney",
"Kidney Cortex",
"Killer Cells  Natural",
"Islets of Langerhans",
"Intestine  Small",
"Intestines",
"Intestinal Mucosa",
"Macrophages",
"Lymphocytes  Null",
"Lymphocytes",
"Lymphoid Tissue",
"Lymph Nodes",
"Lung",
"Liver",
"Limbic System",
"Leukocytes  Mononuclear",
"Leukocytes",
"Mesencephalon",
"Membranes",
"Neutrophils",
"Oocytes",
"Osteoblasts",
"Mouth",
"Mouth Mucosa",
"Monocytes",
"Myometrium",
"Mucous Membrane",
"Muscle  Smooth",
"Muscles",
"Nasopharynx",
"Nasal Mucosa",
"Plasma Cells",
"Ovary",
"Pancreas",
"Parietal Lobe",
"Parotid Gland",
"Periodontium",
"Phagocytes",
"Pharynx",
"Mononuclear Phagocyte System",
"Retina",
"Rectum",
"Prostate",
"Skin",
"Stomach",
"Spleen",
"Stem Cells",
"Salivary Glands",
"Colon  Sigmoid",
"Serous Membrane",
"Thyroid Gland",
"Palatine Tonsil",
"Tongue",
"Testis",
"T Lymphocytes",
"Synovial Membrane",
"Synovial Fluid",
"Vulva",
"Visual Cortex",
"Umbilical Veins",
"Uterus",
"Veins",
"Keratinocytes",
"CD4 Positive T Lymphocytes",
"Adipocytes",
"Stromal Cells",
"Macrophages  Alveolar",
"Chondrocytes",
"Entorhinal Cortex",
"Hepatocytes",
"Myeloid Progenitor Cells",
"Metencephalon",
"Atrial Appendage",
"Mesenchymal Stem Cells",
"Serum",
"Subcutaneous Fat",
"Myocytes  Smooth Muscle",
"Erythroid Cells",
"Upper Gastrointestinal Tract",
"Granulocyte Precursor Cells",
"Gastrointestinal Tract",
"Endothelial Cells",
"Embryonic Stem Cells",
"Foreskin",
"Precursor Cells  B Lymphoid",
"Adult Stem Cells",
"Glucagon Secreting Cells",
"T Lymphocytes  Regulatory",
"Subcutaneous Fat  Abdominal",
"Quadriceps Muscle",
"Neural Stem Cells",
"Embryoid Bodies",
"Induced Pluripotent Stem Cells",
"Granulocyte Macrophage Progenitor Cells",
"Megakaryocyte Erythroid Progenitor Cells",
"Monocyte Macrophage Precursor Cells",
"Endocrine Cells"
)

### heritability enrichment of significantly enriched (FDR < .05) DEPICT gene sets

# Go to
# https://www.gencodegenes.org/human/release_44lift37.html
# for the latest version of this file.
gencode <- read.delim("gencode.v44lift37.basic.annotation.gtf.gz", skip=5, header=FALSE)
# If this file is not present in data_dir, specify the path.
                      
colnames(gencode) <- c("CHR", "SOURCE", "FEATURE", "START", "STOP", "SCORE", "STRAND", "PHASE",
                        "MORE_INFO")  
table(gencode$FEATURE)   

# Create a separate column for Ensembl ID. That makes things much easier.

x <- strsplit(gencode$MORE_INFO, " ")
y <- unlist(lapply(x, function(lst_el) lst_el[2]))
z <- unlist(lapply(strsplit(y, "\\."), function(lst_el) lst_el[1] ))

gencode <- data.frame(gencode, z)
colnames(gencode)[10] <- "ensembl_id"


### significant gene sets for neuroticism general factor
# site of polarized growth GO:0030427
# growth cone GO:0030426
# abnormed cued conditioning behavior MP:0001454
# impaired coordination MP:0001405
# abnormal neuron physiology MP:0004811
# increased anxiety-related response MP:0001363
# decreased exploration in new environment MP:0001417

geneset_matrix_cols <- read.table(paste(
"/Volumes/Backup_Plus/bioinformatics/DEPICT_OLD/data/reconstituted_genesets/",
"GPL570-GPL96-GPL1261-GPL1355TermGeneZScores-MGI_MF_CC_RT_IW_BP_KEGG_z_z.txt.gz", sep=""),
nrows=1
)
# This file is no longer part of the DEPICT download. It is about 2.2G in size. If you would
# like a copy, please email me or the DEPICT developers. My copy probably
# uses too much numerical precision.

geneset_matrix_cols <- as.vector(as.character(geneset_matrix_cols))

which(geneset_matrix_cols=="GO:0030427") # 3507
which(geneset_matrix_cols=="GO:0030426") # 3509
which(geneset_matrix_cols=="MP:0001454") # 1585
which(geneset_matrix_cols=="MP:0001405") # 1119
which(geneset_matrix_cols=="MP:0004811") # 600
which(geneset_matrix_cols=="MP:0001363") # 748
which(geneset_matrix_cols=="MP:0001417") # 1389

# cut these columns as well as the first column (Ensembl IDs) out of the geneset matrix

command <- paste(
"gzcat GPL570-GPL96-GPL1261-GPL1355TermGeneZScores-MGI_MF_CC_RT_IW_BP_KEGG_z_z.txt.gz | ",
"cut -f1,3507,3509,1585,1119,600,748,1389 ",
"> ", data_dir, "neuroticism_general_sig_genesets.txt", sep="")

d_geneset <- read.delim("neuroticism_general_sig_genesets.txt")
colnames(d_geneset) <- c("ensembl_id",
"abnormal_neuron_physiology",
"increased_anxiety_related_response",
"impaired_coordination",
"decreased_exploration_in_new_environment",
"abnormal_cued_conditioning_behavior",
"site_of_polarized_growth",
"growth_cone"
)

out <- subset(gencode, FEATURE=="gene")[,c("ensembl_id", "CHR", "START", "STOP")]
CHR <- unlist(lapply(strsplit(out$CHR, "r"), function(x) x[2]))
out[,"CHR"] <- CHR
colnames(out)[1] <- "GENE"
colnames(out)[4] <- "END"
out <- subset(out, CHR %in% 1:22)
write.table(out, "gencode_coord_file.txt", quote=FALSE, row.names=FALSE, sep="\t")

for(i in 2:ncol(d_geneset)){
	name <- colnames(d_geneset)[i]	
	print(name)
	pct90 <- quantile(d_geneset[,i], 0.9)
	genelist <- d_geneset[d_geneset[,i] > pct90,1]
	write.table(genelist, paste(name, ".GeneSet", sep=""), quote=FALSE, row.names=FALSE)
	
	
for(j in 1:22){	
command <- paste("python ", ldsc_path, "make_annot.py",
                 " --gene-set-file ", data_dir, name, ".GeneSet",
                 " --gene-coord-file ", data_dir, "gencode_coord_file.txt",
                 " --windowsize 100000",
                 " --bimfile ", ldsc_path, "files/1000G_EUR_Phase3_plink/1000G.EUR.QC.", j, 
                 ".bim",
                 " --annot-file ", data_dir,
                 name, ".annot.gz",
                 sep="")
                
                
command <- paste("python ", ldsc_path, "ldsc.py",
                 " --l2",
                 " --bfile ", ldsc_path, "files/1000G_EUR_Phase3_plink/1000G.EUR.QC.", j,
                 " --ld-wind-cm 1",	            
                 " --annot ", data_dir, name, ".", j,
                 ".annot.gz",
                 " --thin-annot",
                 " --out ", data_dir, name, ".", j,
                 " --print-snps ", ldsc_path, "files/1000G_EUR_Phase3_baseline/print_snps.txt",
                 sep="") } # close the loop over chromosomes
                 
command <- paste(ldsc_path, "ldsc.py",
                 " --h2 ", data_dir, "GWASofN_hierarchical.sumstats.gz",
                 " --ref-ld-chr ", data_dir, "reconstituted_", 
                 name, ".",
                 ldsc_path, "files/1000G_Phase3_Franke/Franke.control.,",  # any-gene control              
                 ldsc_path, "files/LDSCORE_1000G_Phase3_baselineLD_v2.2_ldscores/baselineLD.",
                 " --w-ld-chr ", ldsc_path, "files/weights_hm3_no_hla/weights.",
                 " --overlap-annot",
                 " --frqfile-chr ", ldsc_path, "files/1000G_Phase3_frq/1000G.EUR.QC.",
                 " --print-coefficients",
                 " --out ", data_dir, name,
                 sep="") } # close the loop over gene-set names
                 
# Maybe a plot of all S-LDSC enrichments.

geneset_names <- c(
"abnormal_neuron_physiology",
"increased_anxiety_related_response",
"impaired_coordination",
"decreased_exploration_in_new_environment",
"abnormal_cued_conditioning_behavior",
"site_of_polarized_growth",
"growth_cone"
)

geneset_SLDSC_results <- read.delim("reconstituted_abnormal_neuron_physiology.results", nrows=1)
for(i in 2:length(geneset_names)){
	k <- geneset_names[i]
	x <- read.delim(paste("reconstituted_", k, ".results", sep=""), nrows=1)
	geneset_SLDSC_results <- rbind(geneset_SLDSC_results, x) }
	
geneset_SLDSC_results[,1] <- geneset_names

# How do I arrange the results for plotting? 
# Gene set and then tissues. Tissues are in the same order as in the DEPICT tissue plot, except
# arranged within first-level terms by enrichment.

geneset_SLDSC_results <- geneset_SLDSC_results[order(-geneset_SLDSC_results$Enrichment),]

tissue_DEPICT_results <- read.delim(
"../DEPICT_run8/Neuroticism_model_3fact_GWASofN_AIC1_1E-5_tissueenrichment_nodupFranke.txt")

tissue_DEPICT_results <- tissue_DEPICT_results[order(tissue_DEPICT_results$MeSH.first.level.term),]

tissue_SLDSC_results <- d
tissue_SLDSC_results[,1] <- keep
colnames(tissue_SLDSC_results)[1] <- "Name"

SLDSC_results <- merge(tissue_SLDSC_results, 
tissue_DEPICT_results[,c("Name", "MeSH.first.level.term")], by="Name")

SLDSC_results <- SLDSC_results[order(SLDSC_results$MeSH.first.level.term,
                               -SLDSC_results$Enrichment),]
colnames(geneset_SLDSC_results)[1] <- "Name"                               
SLDSC_results <- rbind(geneset_SLDSC_results, SLDSC_results[,1:10])

write.table(SLDSC_results, "all_SLDSC_results.txt", quote=FALSE, sep="\t", row.names=FALSE)

MeSH_first_level_unique <- unique(tissue_DEPICT_results$MeSH.first.level.term)

col_to_plot <- c()
for(i in 1:length(MeSH_first_level_unique)){
	num_tissues <- nrow(subset(tissue_DEPICT_results,
	                MeSH.first.level.term==MeSH_first_level_unique[i]))
	col_to_plot <- c(col_to_plot, rep(brewer.pal(12, "Set3")[i], num_tissues))   }
col_to_plot[138:152] <- "red"	

col_to_plot <- c(rep("black", 7), col_to_plot)
# the third group of colors is too light
col_to_plot[17:51] <- "purple"

quartz(width=8.5, height=5) 

SEfact <- 1

legend_terms <- c(
"GO and MP gene sets",
"Cardiovascular",
"Cells",
"Digestive",
"Endocrine",
"Hemic and immune",
"Integumentary",
"Musculoskeletal",
"Nervous",
"Respiratory",
"Sense organs",
"Stomatognathic",
"Tissues",
"Urogenital"
)

plot(1:159, rep(0, 159), col="grey60", pch=17, ylim=c(0, 2), xlab="",
     ylab="Heritability enrichment", axes=FALSE, type="n")

# error bars have to be first so that data are "on top"

arrows(x0=1:159, y0=SLDSC_results$Enrichment - SEfact*SLDSC_results$Enrichment_std_err,
                 y1=SLDSC_results$Enrichment + SEfact*SLDSC_results$Enrichment_std_err,
                 length=0, lwd=0.5, col=col_to_plot)

points(1:7, SLDSC_results$Enrichment[1:7], col=col_to_plot[1:7], pch=17, cex=2/3)
points(8:159, SLDSC_results$Enrichment[8:159], col=col_to_plot[8:159], pch=19, cex=0.5)

legend(x="bottomright", legend=legend_terms, fill=unique(col_to_plot), cex=0.75,
       ncol=3, text.width=30, bty="n")
     
axis(2, at=seq(0, 2, 0.25))
abline(h=1, lwd=0.5)
abline(h=1.3, lwd=0.5, lty="dashed")

segments(2, 1.48, 2, 1.75)
segments(2, 1.75, 5, 1.75)
text(4.25, 1.75, pos=4, labels="Increased anxiety-related response", cex=0.75)

segments(3, 1.42, 3, 1.6)
segments(3, 1.6, 6, 1.6)
text(5, 1.6, pos=4, labels="Decreased exploration in new environment", cex=0.75)

segments(4, 1.41, 4, 1.45)
segments(4, 1.45, 7, 1.45)
text(6, 1.45, pos=4, labels="Abnormal cued conditioning behavior", cex=0.75)

segments(17, 1.22, 17, 1.15)
segments(17, 1.15, 20, 1.15)
text(19, 1.15, pos=4, labels="Neural stem cells", cex=0.75)

segments(121, 1.40, 121, 1.46)
segments(121, 1.46, 125, 1.46)
text(124, 1.46, pos=4, labels="Retina", cex=0.75)  

segments(102, 1.51, 102, 1.61) 
segments(102, 1.61, 105, 1.61)                                              
text(104, 1.61, pos=4, labels="Entorhinal cortex", cex=0.75)

dev.off()
```
